# Structural remodeling of the mitochondrial protein biogenesis machinery under proteostatic stress

**DOI:** 10.1101/2025.10.06.680693

**Authors:** Kenneth Ehses, Jorge P. López-Alonso, Odetta Antico, Abdussalam Azem, Miratul M.K. Muqit, Iban Ubarretxena-Belandia, Rubén Fernández-Busnadiego

## Abstract

Cells have evolved organelle-specific responses to maintain protein homeostasis (proteostasis). During proteostatic stress, mitochondria downregulate translation and enhance protein folding, yet the underlying mechanisms remain poorly defined. Here, we employed cryo-electron tomography to observe the structural consequences of mitochondrial proteostatic stress within human cells. We detected protein aggregates within the mitochondrial matrix, accompanied by a marked remodeling of cristae architecture. Concomitantly, the number of mitochondrial ribosome complexes was significantly reduced. Mitochondrial Hsp60 (mHsp60), a key protein folding machine, underwent major conformational changes to favor complexes with its co-chaperone mHsp10. We visualized the interactions of mHsp60 with native substrate proteins, and determined *in vitro* mHsp60 cryo-EM structures enabling nucleotide state assignment of the *in situ* structures. These data converge on a model of the mHsp60 functional cycle and its essential role in mitochondrial proteostasis. More broadly, our findings reveal structural mechanisms governing mitochondrial protein biosynthesis and their remodeling under proteostatic stress.

## Introduction

Most mitochondrial proteins are encoded in the nuclear genome and must be imported into the organelle upon cytosolic translation. For proteins destined to the mitochondrial matrix, import requires threading in an extended conformation through the narrow pores of the translocases of both the outer and inner mitochondrial membranes (TOM and TIM23 complexes, respectively) (den Brave *et al*, 2024; Jain *et al*, 2025). Once in the matrix, these polypeptides must immediately fold into their native conformation. Mitochondria harbor a dedicated protein quality control network, encompassing chaperones of the mitochondrial heat shock protein (mHsp) families mHsp60, mHsp70, mHsp90 and mHsp100/Clp, as well as proteases such as Lonp1, which assist in folding nascent or stress-denatured proteins and degrade irreversibly misfolded polypeptides, respectively (Song *et al*, 2021; Voos *et al*, 2016).

The mHsp60:mHsp10 chaperonin is a central protein folding machine in this system (Cheng *et al*, 1989; Ostermann *et al*, 1989; Reading *et al*, 1989). mHsp60:mHsp10 assists the folding of approximately half of all matrix proteins, and corrects polypeptides misfolded under stress (Bender *et al*, 2011; Bie *et al*, 2020). Reflecting its essential role, deletion or inactivation of the mHsp60 gene is lethal in yeast and mammals (Cheng *et al*., 1989; Christensen *et al*, 2010), and mutations in the mHsp60 gene lead to severe neurodegenerative diseases, including hereditary spastic paraplegia and hypomyelinating leukodystrophy (Bross & Fernandez-Guerra, 2016; Hansen *et al*, 2002; Magen *et al*, 2008). Structurally, mHsp60 harbors an equatorial domain responsible for nucleotide binding, an intermediate domain with two hinge regions that facilitate conformational changes, as well as an apical domain involved in binding substrate proteins (SPs) and the mHsp10 protein cofactor (Nisemblat *et al*, 2015). mHsp60 protomers assemble into heptameric rings, which can be capped by heptameric dome-shaped mHsp10 lids to form nanocages for SPs to fold in confinement (Gomez-Llorente *et al*, 2020; Nisemblat *et al*., 2015; Weiss *et al*, 2016). Although much of our current understanding of mHsp60:mHsp10 function is derived from the highly homologous bacterial GroEL:GroES system (Hayer-Hartl *et al*, 2016; Horwich & Fenton, 2020), biochemical and structural evidence indicates key differences between the two (Gomez-Llorente *et al*., 2020; Ishida *et al*, 2018; Okamoto *et al*, 2015; Weiss *et al*., 2016). Thus, the mechanisms of protein folding by mHsp60 warrant further investigation.

Mitochondria also possess their own genome, which in mammals codes for 13 proteins, 11 messenger RNAs, 22 transfer RNAs and 2 ribosomal RNAs (Kummer & Ban, 2021). Dedicated molecular machines carry out the transcription and translation of this genome, including a mitochondrial ribosome complex with substantial structural and functional differences compared to its cytosolic counterpart (Kummer & Ban, 2021; Vuckovic *et al*, 2024). Nuclear- and mitochondrial-encoded proteins are co-assembled into large complexes of the oxidative phosphorylation machinery, and nuclear-encoded chaperones and proteases are also essential for the folding and degradation of mitochondrially-translated polypeptides (Kummer & Ban, 2021; Song *et al*., 2021). Therefore, mitochondrial protein biosynthesis depends on an intricate coordination between nuclear- and mitochondrial-encoded factors, many aspects of which remain only partly understood.

Disruptions in mitochondrial proteostasis can activate protective cellular pathways, most notably the so-called mitochondrial unfolded protein response (UPR^mt^). UPR^mt^ is triggered amongst others by the accumulation of misfolded proteins in the mitochondrial matrix (Munch, 2018; Shpilka & Haynes, 2018), and involves both local mitochondrial responses and nuclear transcription programs. This profoundly affects the mitochondrial protein biosynthesis machinery by acutely downregulating protein translation and enhancing the expression of protein folding factors. Among the latter, mHsp60 and mHsp10 are considered canonical UPR^mt^ responders (Martinus *et al*, 1996; Munch & Harper, 2016; Yoneda *et al*, 2004; Zhao *et al*, 2002). Chronic failure to restore the fine balance of mitochondrial proteostasis can lead to pathological conditions, with proteostatic deficits associated to neurodegenerative diseases such as Alzheimer’s and Parkinson’s (Chen *et al*, 2023; Eldeeb *et al*, 2022; Sorrentino *et al*, 2017), and excessive capacity linked to tumor growth and metastasis (Inigo & Chandra, 2022; Zhang *et al*, 2024). Thus, it is essential to gain detailed understanding of the molecular and structural mechanisms that safeguard mitochondrial protein homeostasis.

Here, we tackle this challenge using cryo-electron tomography (cryo-ET), which enables 3D visualization of cellular machinery under close-to-native conditions within vitrified cells (Bauerlein & Baumeister, 2021; Nogales & Mahamid, 2024). We have previously employed cryo-ET to analyze the effects of proteostatic challenges such as cytosolic protein aggregation (Bauerlein *et al*, 2017; Gruber *et al*, 2018; Guo *et al*, 2018; Riemenschneider *et al*, 2022; Riera-Tur *et al*, 2022; Trinkaus *et al*, 2021), and to investigate the functional cycle of GroEL:GroES in *E.coli* cells (Wagner *et al*, 2024). Here, we capitalize on cryo-ET to study the structural consequences of proteostatic stress in mitochondria by imaging these organelles within human cells at subnanometer resolution. Our data reveal the formation of protein aggregates in the mitochondrial matrix, accompanied by major morphological rearrangements of the mitochondrial inner membrane, as well as remodeling of the conformational landscape of mitochondrial ribosomes and mHsp60:mHsp10 complexes. Analysis of mHsp60-SP interactions *in situ* combined with single-particle cryo-EM imaging of purified proteins converge on a working model for the mHsp60:mHsp10 functional cycle in cells and its adaptation to misfolding stress. Collectively, this work provides structural insights of how mitochondria remodel their biosynthetic machinery under proteostatic stress.

## Results

### Mitochondria morphological remodeling under proteostatic stress

Under proteostatic stress, damaged mitochondria may be cleared by PINK1-Parkin mediated mitophagy (Fiesel *et al*, 2017; Jin & Youle, 2013; Michaelis *et al*, 2022; Uoselis *et al*, 2023), complicating their analysis by cryo-ET. To circumvent this issue, we studied mitochondrial proteostatic stress in HeLa cells, which naturally lack Parkin and thus display low mitophagy levels (Burman *et al*, 2017; Denison *et al*, 2003). HeLa cells were genetically edited by CRISPR-Cas9 to introduce an endogenous C-terminal GFP tag at the PINK1 locus, enabling facile monitoring of mitochondrial stress by imaging the mitochondrial accumulation of PINK1 (Narendra & Youle, 2024; Trempe & Gehring, 2023). We induced mitochondrial proteostatic stress by treating cells with gamitrinib-triphenylphosphonium (G-TPP), a specific inhibitor of the mHsp90 chaperone TRAP1 that causes intramitochondrial protein aggregation, PINK1 stabilization on mitochondria and UPR^mt^ activation (Fiesel *et al*., 2017; Kang *et al*, 2009; Michaelis *et al*., 2022; Munch & Harper, 2016; Sutandy *et al*, 2023; Uoselis *et al*., 2023). Consistent with the induction of mitochondrial stress, PINK1-GFP imaging and western blot analysis revealed robust PINK1 activation on mitochondria following 4 h G-TPP treatment (Figure S 1A, C). Cell viability analyses showed that, whereas no significant changes appeared upon a 4 h G-TPP treatment, cell health began to significantly deteriorate beyond 6 h (Figure S 1B). Thus, our cryo-ET imaging focused on the 4 h time point, where proteostatic responses have been activated but no major confounding effects from cell death should be present. To that end, HeLa cells were cultured on EM grids, treated with G-TPP for 4 h (“Proteostatic stress” condition) or left untreated (“Control” condition), vitrified by plunge-freezing and thinned down to ∼150 nm-thick lamellae using cryo-focused ion beam milling (cryo-FIB; Figure S 1D). Mitochondria were visually located on the resulting lamellae (Figure S 1E) and imaged by cryo-ET. This revealed no overt changes in cellular architecture upon 4 h G-TPP treatment (Figure 1A, B; Figure S 1F, G), consistent with our cell viability measurements (Figure S 1B). However, under these conditions ∼80% of mitochondria (N=137; Figure 1B; Figure S 1E, G) contained dense accumulations within their matrix, likely corresponding to protein aggregates (Kang *et al*., 2009; Uoselis *et al*., 2023), which appeared amorphous at our resolution. These aggregates were not observed in any control mitochondria (N=105; Figure 1A; Figure S 1F). Thus, our experimental setup allowed cryo-ET investigations of the structural consequences of mitochondrial proteostatic stress within native cellular environments.

**Figure 1:**
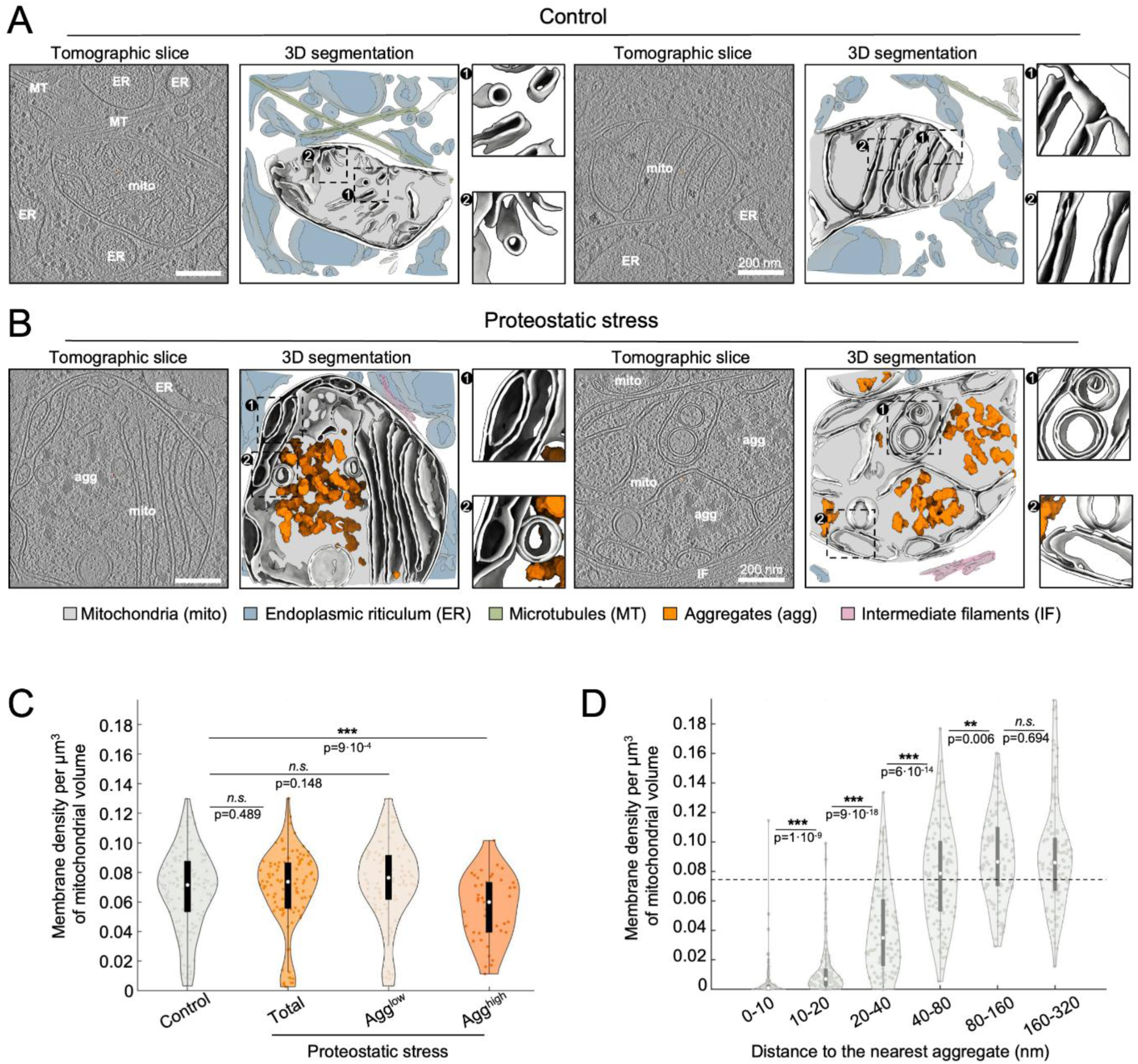
Proteostatic stress triggers cristae remodeling and aggregate formation within mitochondria. Representative tomographic slices and 3D segmentations from untreated (“Control”) (**A**) and G-TPP-treated (“Proteostatic stress”) HeLa cells (**B**). In the 3D segmentations, mitochondria are depicted in grey and additional cellular elements according to the color code in the figure. (**C**) Quantification of cristae density within mitochondria. For the proteostatic stress condition, data is shown for all cells, and also separated by aggregate load into Agg^low^ (mitochondria with low aggregate load or no aggregates) and Agg^high^ (mitochondria with high aggregate load). (**D**) Cristae density as a function of the distance to the nearest aggregate. Statistical significance of pairwise comparisons in (C) and (D) was assessed using a nonparametric Wilcoxon rank-sum test and indicated by: n.s. (p > 0.05), * (p < 0.05), ** (p < 0.01) and *** (p < 0.001).

At the morphological level, we observed substantial cristae remodeling under proteostatic stress. In control mitochondria, cristae were generally planar and arranged parallel to each other, with cristae junctions forming near 90-degree angles with the outer mitochondrial membrane (Figure 1A; Figure S 1F). In contrast, upon proteostatic stress we observed numerous cristae at near-parallel angles to the outer mitochondrial membrane, as well as circular cristae (Figure 1B; Figure S 1G). Whereas proteostatic stress did not lead to global changes in cristae density, this parameter was significantly reduced in the subgroup of mitochondria with higher aggregate load (Figure 1C). In most cases, cristae alterations appeared local, with a reduction in cristae density in the immediate vicinity of aggregates, but not in more distal regions (Figure 1B left, Figure 1D; Figure S 1G). However, other mitochondria showed generalized cristae alterations (Figure 1B right; Figure S 1G), potentially corresponding to a more advanced stage of damage. These data suggest that upon proteostatic stress, aggregate formation within the mitochondrial matrix leads to local cristae deformations and eventually their disappearance. This phenomenon may extend throughout the entire organelle when stress persists, resulting in a major disruption of cristae organization.

### Proteostatic stress reconfigures the mitochondrial biogenesis machinery

We investigated how proteostatic stress may impact the mitochondrial machineries for protein synthesis and folding. We first focused on mitochondrial ribosomes, as G-TPP treatment has been reported to reduce mitochondrial protein translation and cause the sequestration of mitochondrial ribosome subunits into protein aggregates (Holthusen *et al*, 2025; Munch & Harper, 2016; Uoselis *et al*., 2023). We located mitochondrial ribosomes in cells under control and stress conditions using template matching (Figure S 2A, B), and these initial hits were used to generate a data-derived template for further extensive template matching searches (Figure S 2A, B). Upon subsequent alignment, averaging and classification (Figure S 3A, B), this process converged to two structures, corresponding to the 39S large ribosomal subunit and the fully assembled 55S mitochondrial ribosome (Figure 2A). The structures were determined at ∼16 Å and ∼23 Å nominal resolution, respectively (Figure S 3C), and showed a prominent density for the membrane bilayer (Figure 2A, Figure S 3C), indicating that most ribosomal complexes detected were anchored to the inner mitochondrial membrane. Available atomic models derived from published single-particle structures of the 39S and 55S mitochondrial ribosomes fitted well our subtomogram averaging maps (Figure S 4A, B). Of note, our 39S map displayed a density consistent with the anti-association module comprising MALSU1, L0R8F8 and mtACP (Figure S 4A), which has been reported to block premature small subunit binding during ribosome assembly (Hillen *et al*, 2021; Itoh *et al*, 2022; Lavdovskaia *et al*, 2024; Rebelo-Guiomar *et al*, 2022). In contrast, the 55S mitochondrial ribosome displayed clear density for the 28S small ribosomal subunit (Figure S 4B), along with additional densities consistent with components found exclusively in translation-competent mitochondrial ribosomes, such as the RNA-binding protein PTCD3/mS39 (Kummer *et al*, 2018). Notably, proteostatic stress triggered a highly significant reduction of both 39S and 55S ribosomal complexes, which was even more pronounced for the latter (Figure 2B, C). Thus, our data indicate that proteostatic stress strongly reduces the number of translation-competent mitochondrial ribosome complexes.

**Figure 2:**
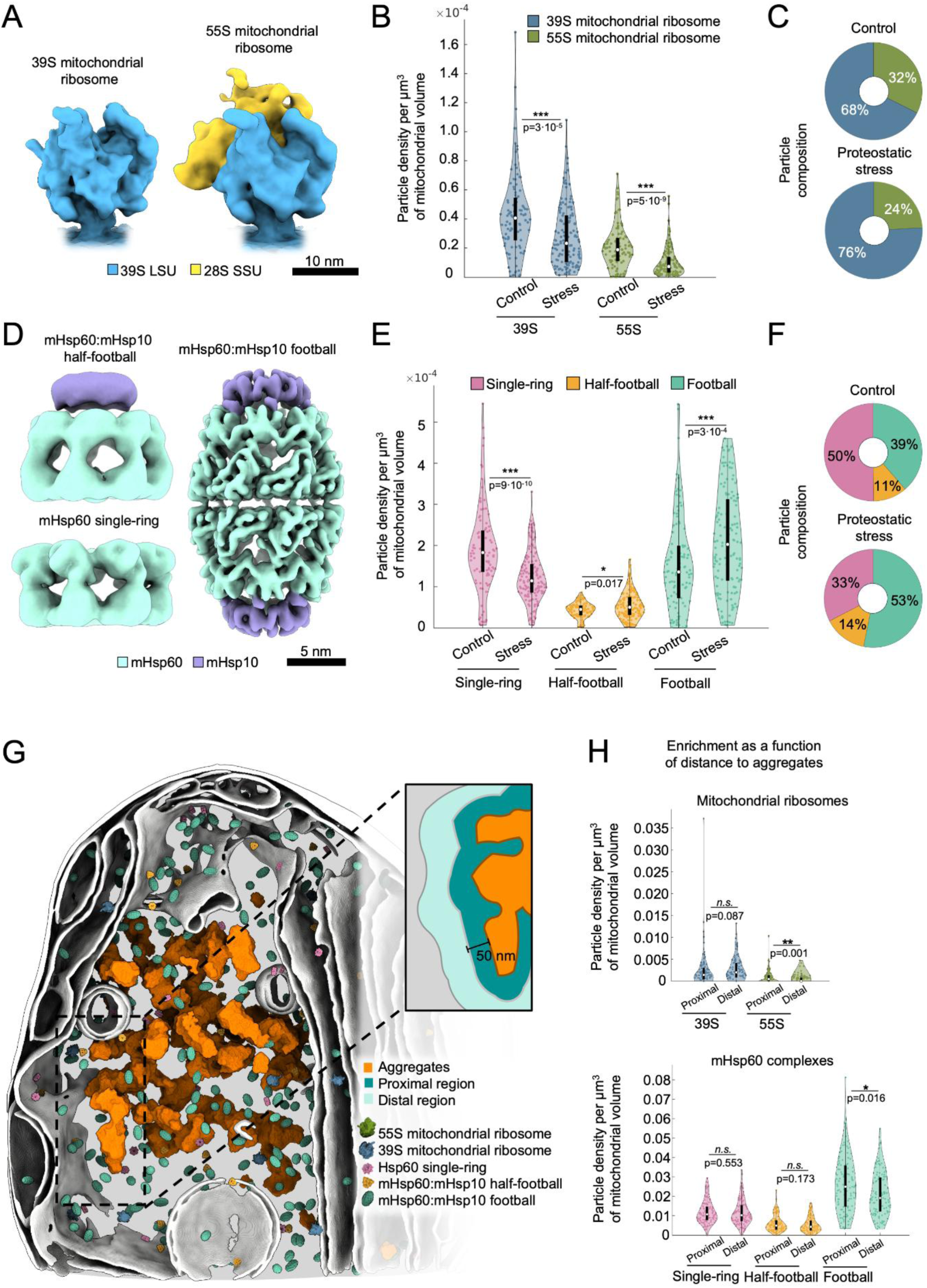
*In situ* structures reveal the native structural landscape of mitochondrial ribosomes and mHsp60:mHsp10 complexes and its reorganization under proteostatic **stress.** (**A**) Subtomogram averages of the 39S large mitochondrial subunit and fully assembled 55S mitochondrial ribosome. (**B**) Spatial density of mitochondrial ribosome complexes in control and proteostatic stress conditions. (**C**) Relative abundance of mitochondrial ribosomal complexes. (**D**) Subtomogram averages of mHsp60 complexes. (**E**) Density of mHsp60 complexes in control and proteostatic stress conditions. (**F**) Relative abundance of mHsp60 complexes. (**G**) 3D visualization of mitochondrial ribosomes and mHsp60 complexes within an aggregate-containing mitochondrion in a cell under proteostatic stress. The inset illustrates the definitions of the “proximal” and “distal” areas relative to the aggregates used in the quantifications shown in (H). The proximal area encompasses particles up to 50 nm away from the aggregates. The distal area extends further than the proximal area until an equivalent volume is reached. (**H**) Spatial distribution of ribosomal (top) and mHsp60 (bottom) complexes in cells under proteostatic stress as a function of the distance to the aggregate. Statistical significance of pairwise comparisons in (B), (E) and (H) was assessed using a nonparametric Wilcoxon rank-sum test and indicated by: n.s. (p > 0.05), * (p < 0.05), ** (p < 0.01) and *** (p < 0.001).

We next focused on the mHsp60:mHsp10 chaperonin system, a major responder to mitochondrial proteostatic stress and canonical UPR^mt^ effector (Martinus *et al*., 1996; Munch & Harper, 2016; Yoneda *et al*., 2004; Zhao *et al*., 2002). Multiple mHsp60 oligomeric species have been observed *in vitro*, including mHsp60 single-rings (mHsp607) and mHsp60 double rings (mHsp6014), as well as various complexes with its protein cofactor mHsp10, such as mHsp60:mHsp10 half-football (mHsp607:mHsp107), bullet (mHsp6014:mHsp107) and football complexes (mHsp6014:mHsp1014) (Braxton *et al*, 2024; Gomez-Llorente *et al*., 2020; Klebl *et al*, 2021; Levy-Rimler *et al*, 2001; Nisemblat *et al*., 2015; Okamoto *et al*., 2015; Syed *et al*, 2024). However, it remains uncertain which of these assemblies participate directly in the mHsp60 SP folding cycle within mitochondria. Thus, we searched our tomograms for all the above-mentioned mHsp60 and mHsp60:mHsp10 complexes using template matching by cropping the relevant portions (Figure S 2A, C-G) of our published cryo-EM structure of the mHsp60:mHsp10 football (Gomez-Llorente *et al*., 2020). This search converged for mHsp60 single-rings, as well as for mHsp60:mHsp10 half-footballs and footballs, but not for mHsp60 double rings and mHsp60:mHsp10 bullets. These results indicate that in mitochondria of HeLa cells, the predominant mHsp60 oligomeric species are single-rings, half-footballs and footballs.

Following an analogous procedure to that used for mitochondrial ribosomes, we determined structures of the detected mHsp60 species using subtomogram averaging (Figure S 3D, E). This resulted in maps at ∼14 Å, ∼14 Å and ∼8 Å nominal resolution for the mHsp60 single-rings, mHsp60:mHsp10 half-footballs and mHsp60:mHsp10 footballs, respectively (Figure 2D, Figure S 3F). In control cells, mHsp60 single-rings were the most abundant species, followed by mHsp60:mHsp10 footballs, with mHsp60:mHsp10 half-footballs constituting a minor fraction (Figure 2E, F). Proteostatic stress led to a substantial rearrangement of this assembly landscape, favoring mHsp60:mHsp10 complexes, especially mHsp60:mHsp10 footballs, and strongly reducing mHsp60 single-rings (Figure 2E, F). Therefore, protein aggregation favored the assembly of mHsp60:mHsp10 complexes, in particular football complexes containing two folding chambers.

We then analyzed the distribution of the ribosomal and mHsp60 species detected as a function of their distance to the aggregates by defining “proximal” and “distal” regions of equivalent volume (Figure 2G). 39S large ribosomal subunits, mHsp60 single-rings and mHsp60:mHsp10 half-football complexes did not show a preferred distribution across these regions (Figure 2H). However, 55S ribosomes were depleted within the proximal region (Figure 2H), consistent with a direct effect of protein aggregation in the reduction of mitochondrial translation upon proteostatic stress (Holthusen *et al*., 2025; Munch & Harper, 2016; Uoselis *et al*., 2023). In contrast, mHsp60:mHsp10 footballs were enriched in the proximal region (Figure 2H), pointing to the preferential formation of these complexes under high abundance of misfolded SPs. These data suggest that protein aggregate formation may play a direct role in the structural and spatial reorganization of the mitochondrial biosynthesis machinery observed under proteostatic stress.

### mHsp60 single-rings in mitochondria are bound to ADP

Available high-resolution structures from us and others, together with molecular dynamics simulations, indicate that ATP binding and hydrolysis drive the conformational and assembly dynamics of mHsp60:mHsp10 complexes during their SP folding cycle (Braxton *et al*., 2024; Gomez-Llorente *et al*., 2020; Klebl *et al*., 2021; Nisemblat *et al*., 2015; Syed *et al*., 2024; Tascon *et al*, 2025; Torielli *et al*, 2025; Wang & Chen, 2021). Therefore, to understand the functional status of the mHsp60 complexes detected in our *in situ* tomograms (Figure 2D), it is important to gain insight into their nucleotide state. However, the resolution of our subtomogram averaging maps did not allow direct identification of the nucleotides. At the same time, ambiguities persisted upon fitting the high-resolution structures available (Figure S 4C- F), as reported structures of wild-type (WT) and mutant mHsp60:mHsp10 complexes in different nucleotide states (Braxton *et al*., 2024; Gomez-Llorente *et al*., 2020; Tascon *et al*., 2025) fitted our half-football and football maps (Figure S 4E, F; Table S 1). Furthermore, neither of the available structures of apo or ATP-bound mHsp60 single-rings (Braxton *et al*., 2024; Syed *et al*., 2024; Tascon *et al*., 2025; Wang & Chen, 2021) fitted well the conformation of the mHsp60 apical domains observed *in situ* (Figure S 4C, D), suggesting that the *in situ* single-ring structure corresponded to a conformation not previously observed *in vitro*.

To address these limitations, we carried out single-particle cryo-EM studies of recombinant WT human mHsp60 and mHsp10 complexes. Mixing mHsp60 with mHsp10 in a 0.8:1 ratio in the presence of ATP resulted in cryo-EM maps of the following complexes (Figure 3A, Figure S 5A, B, Table S 2): ATP-bound mHsp60:mHsp10 football (1.92 Å nominal resolution), ATP-bound mHsp60:mHsp10 half-football (2.19 Å nominal resolution), ATP-bound mHsp60:mHsp10 bullet (2.1 Å nominal resolution), ATP-bound mHsp60 double ring (2.18 Å nominal resolution), ADP-bound mHsp60 single-ring (2.91 Å nominal resolution) and ATP-bound mHsp60 single-ring (2.50 Å nominal resolution). The maps compared well with available structures (Table S 1), except for the ATP-bound mHsp60:mHsp10 bullet and the ADP-bound mHsp60 single-ring structures, which were not reported before. These data confirm that all possible combinations of heptameric mHsp60 and mHsp10 assemblies can form *in vitro*, although only mHsp60 single-rings, mHsp60:mHsp10 half-football and football complexes were observed *in situ* within the mitochondria of HeLa cells.

**Figure 3:**
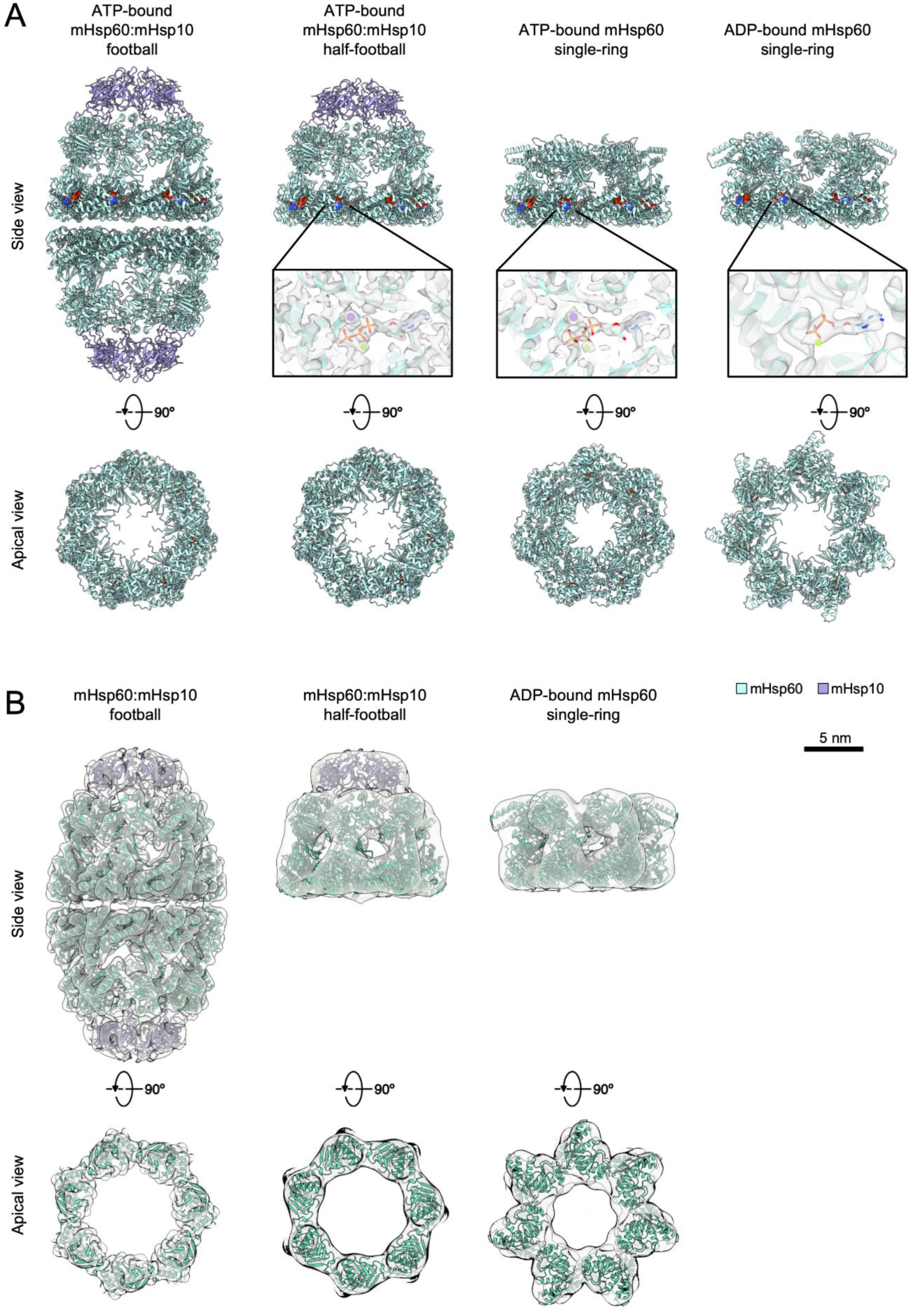
Single-particle cryo-EM structures of purified mHsp60 complexes enable nucleotide state assignment of *in situ* mHps60 assemblies. (**A**) Ribbon representation of the atomic models derived from single-particle cryo-EM maps of the ATP-bound mHsp60:mHsp10 football (mHsp6014:mHsp1014), ATP-bound mHsp60:mHsp10 half-football (mHsp6014:mHsp107), ATP-bound mHsp60 single-ring (mHsp607) and ADP-bound mHsp60 single-ring, showing mHsp60 subunits in cyan and mHsp10 subunits in purple. The models are shown in side view (top) and in a top view omitting the mHsp10 lids to better visualize the mHsp60 apical domains (bottom). Space-filling representations depict the nucleotides in CPK color scheme. Insets show overlays of the density map (semitransparent) and atomic models in the nucleotide binding pockets. Nucleotides are depicted as sticks, and Mg^2+^ and K^+^ ions are shown as green and purple spheres, respectively. (**B**) Rigid-body docking of the single- particle-derived atomic models into the corresponding subtomogram maps obtained *in situ* (semitransparent). Overlays are shown in side view (top) and in top view at the region corresponding to the apical domains (bottom).

We then analyzed in more detail the single-particle cryo-EM structures corresponding to the mHsp60 species visualized in cells. Inspection of the mHsp60:mHsp10 half-football and football cryo-EM density maps revealed the presence of ATP in all the nucleotide-binding pockets, even when no symmetry was applied to the 3D reconstructions (Figure 3A, Figure S 5C). This is consistent with available football structures, where the same nucleotide occupies both rings of the football (Braxton *et al*., 2024; Gomez-Llorente *et al*., 2020; Nisemblat *et al*., 2015; Tascon *et al*., 2025). Atomic models derived from the single-particle maps of ATP-bound mHsp60:mHsp10 half-football and football accurately fitted our respective subtomogram averaging maps (Figure 3B, Figure S 4E, F), leading us to tentatively assign the mHsp60:mHsp10 structures obtained in cells to ATP-bound states.

Next, we investigated in depth the single-particle structures obtained for mHsp60 single-rings. Classification revealed two major nucleotide-binding states, corresponding to ATP- and ADP-bound conformations (Figure 3A, Figure S 5A, B). The ATP-bound mHsp60 single-ring structure was almost identical to that recently described for each ring of the ATP-bound double-ring WT mHsp60 (Tascon *et al*., 2025) and the V72I variant (Braxton *et al*., 2024). These structures did not fit well the apical domains of our *in situ* mHsp60 single-ring map, and displayed additional densities on the central pore (Figure S 4C). In contrast, the ADP-bound mHsp60 single-ring structure accurately fitted the mHsp60 single-ring subtomogram averaging map (Figure 3B, Figure S 4C, D), suggesting that mHsp60 single-rings in cells may largely correspond to the ADP-bound species.

ADP-bound mHsp60 single-rings had not been reported previously, thus warranting detailed analysis of this structure. The most striking difference between the ATP- and ADP-bound mHsp60 single-ring structures is the conformation of the apical domains, which are much better defined in the ADP-bound structure (Figure S 5B). We first analyzed the conformation of helices H and I, responsible for binding SPs and mHsp10 (Braxton *et al*., 2024; Gomez-Llorente *et al*., 2020; Nisemblat *et al*., 2015; Tascon *et al*., 2025). In the ATP-bound mHsp60 single-ring, helices H and I are facing the central cavity of the ring, in a dynamic alternating up/down configuration that has been proposed to help SP capture (Braxton *et al*., 2024). In ATP-bound footballs, the mHsp60 apical domains undergo a vertical rigid body rotation that elevates helices H and I, thereby releasing the SP, enabling mHsp10 binding and SP folding within the mHsp60:mHsp10 chamber (Braxton *et al*., 2024; Gomez-Llorente *et al*., 2020; Nisemblat *et al*., 2015; Tascon *et al*., 2025). However, in the ADP-bound mHsp60 single-rings, the apical domains adopt a different orientation, undergoing a horizontal rigid body rotation relative to the ATP-bound single-ring form (Figure 4A). This positions helices H and I sideways from the central cavity, potentially reducing their capacity to bind both SPs and mHsp10.

**Figure 4:**
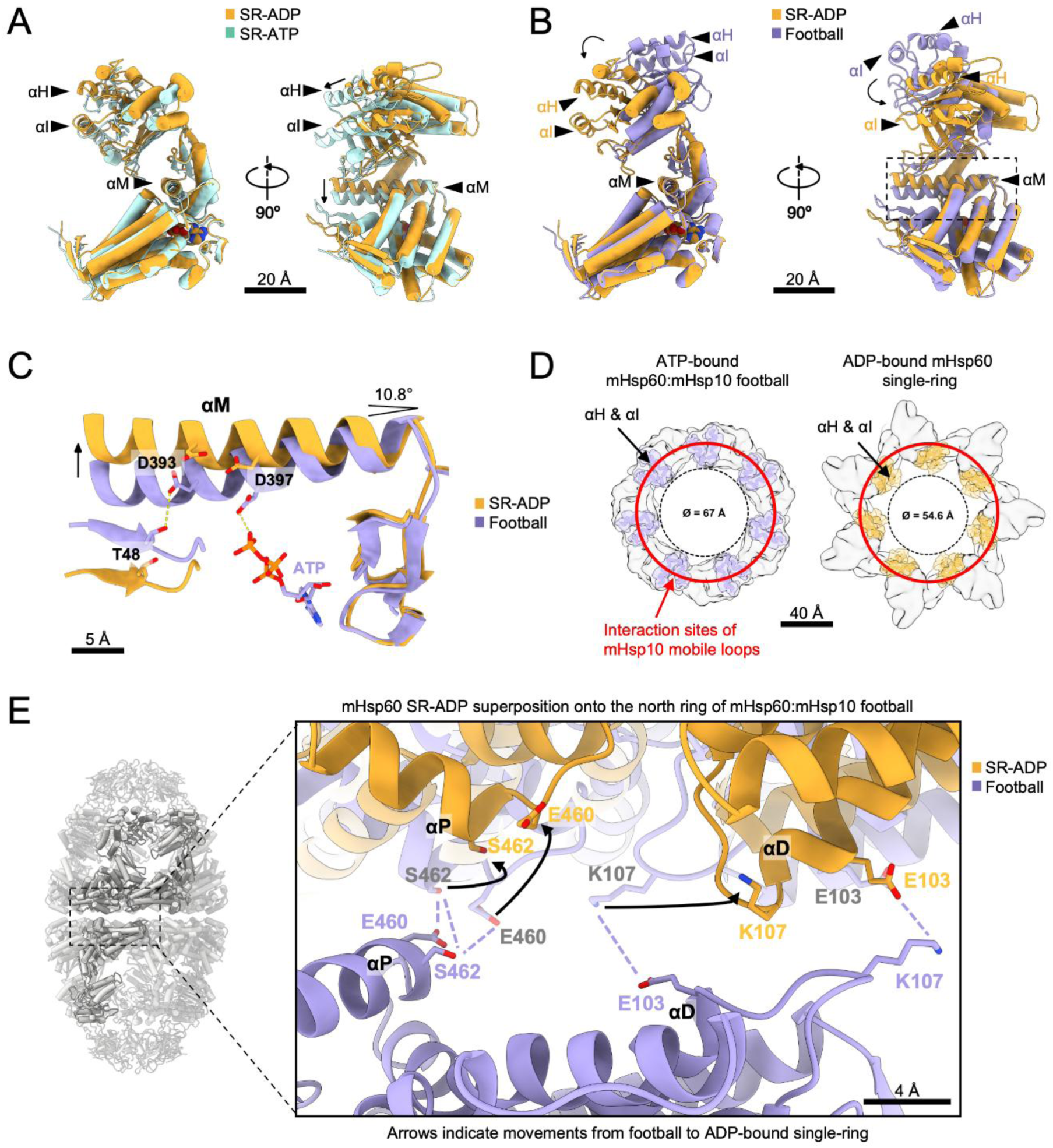
The structure of the ADP-bound mHsp60 single-ring reveals ATP-driven conformational changes in the chaperonin functional cycle. (**A**) Superposition of the mHsp60 monomer in the ATP-bound (cyan) and the ADP-bound (yellow) single-ring (SR) structures. Relative to the ADP-bound single-ring, exchange of ADP by ATP in the mHsp60 single-ring results in a rigid body horizontal rotation of the apical domain (see helices H [αH] and I [αI]) and downward pivoting of helix M (αM) in the intermediate domain. (**B**) Superposition of the mHsp60 monomer in the ATP-bound mHsp60:mHsp10 football (purple) and the ADP-bound mHsp60 single-ring (yellow) structures. The boxed area is magnified in (C). (**C**) Close view of the nucleotide-binding pocket marked by a box in (B) for the mHsp60 monomer in the ATP-bound mHsp60:mHsp10 football (purple) and the ADP-bound mHsp60 single-ring (yellow) structures. This shows how ATP hydrolysis and the loss of the γ-phosphate breaks the contact with residue D397, resulting in the ∼11° upward pivoting of helix M. (**D**) Top cartoon views of the ATP-bound mHsp60:mHsp10 football (purple) and ADP-bound mHsp60 single-ring (yellow) structures showing only the apical domains with highlighted helices H and I. A red circle marks the sites of interaction of the mHsp10 mobile loops with mHsp60 apical helices H and I in the mHsp60:mHsp10 football structure. In the ADP-bound mHsp60 single- ring, the apical helices H and I move away from the mHsp10 landing sites preventing the interaction. Note also the differences in the diameter of the central pore. For clarity, in panels A-C helices H and I in the apical domain and Helix M in the intermediate domain are highlighted as ribbons, whereas cylinders denote the remaining helices of the protein. Arrows denote the direction of movement. (**E**) Close view of the equatorial inter-ring interface in the ATP-bound mHsp60:mHsp10 football (purple) structure with the superimposed ADP-bound mHsp60 single-ring structure (yellow). In the ATP-bound football, interactions between equatorial helices D (αD) and P (αP) stich the two rings together. ATP hydrolysis causes conformational changes that alter the positions of both helices preventing the interactions between E460 and S462, as well asbetween K107 and E103 across the interface. Arrows mark the movement of key residues in the ADP-bound mHsp60 single-ring, which are not compatible with equatorial dimerization.

The analysis of the mHsp60 equatorial and intermediate domains provided a rationale for these observations. Whereas in the ATP-bound football structure D397 on helix M binds to the γ phosphate of ATP, this interaction is lost upon ATP hydrolysis (Figure 4C). This causes helix M to tilt upwards, driving an allosteric conformational change of the apical domain. As this conformation is not compatible with mHsp10 binding (Figure 4B, D), ATP hydrolysis within football and half-football complexes is likely to trigger ejection of mHsp10. Further comparison of the ATP-bound mHsp60:mHsp10 football and the ADP-bound mHsp60 single-ring also revealed differences in the inter-ring interface (Figure 4E). In the ATP-bound mHsp60:mHsp10 football helices D and P form a smooth interface, allowing inter-ring interactions between E460 and S462, as well as E103 and K107 of an adjacent subunit (Gomez-Llorente *et al*., 2020; Tascon *et al*., 2025). In contrast, the equatorial interface of the ADP-bound mHsp60 single-ring structure is incompatible with these inter-ring interactions (Figure 4E). Therefore, our structures suggest that ATP hydrolysis within mHsp60:mHsp10 football complexes leads to both mHsp10 ejection and equatorial split. Thus, the abundant ADP-bound mHsp60 single-rings observed in cells (Figure 2D-F, Figure 3B) could result from the dissociation of mHsp60:mHsp10 complexes upon ATP hydrolysis.

### Interactions between mHsp60 single-rings and substrate proteins *in situ*

To further investigate the functional roles of the mHsp60 single-rings detected in cells, we analyzed their interactions with SPs. We classified mHsp60 single-ring particles using a mask focused on their central cavity (Figure S 6A). This process revealed four different groups: a class where no density was visible on the mHsp60 central cavity, and another three classes where additional densities were present at different heights of the cavity, either only at the equatorial region or both at the apical and equatorial domains (Figure 5A). The classification was robust, as similar classes were obtained upon independent runs (Figure S 6B). However, at the resolution obtained it was not possible to conclude with certainty whether the “no density” class represented the absence of bound SP, dynamic SP interactions that are lost by averaging, or a mixture of both. In any case, the “no density” class was dominant in control cells, and proteostatic stress did not cause major changes in the relative abundance of the different classes (Figure 5C). Therefore, although the relative proportions of mHsp60 single-rings and mHsp60:mHsp10 half-footballs and footballs changed significantly upon stress (Figure 2E, F), proteostatic stress may not affect the dynamics of the interactions between mHsp60 single-rings and SPs.

**Figure 5:**
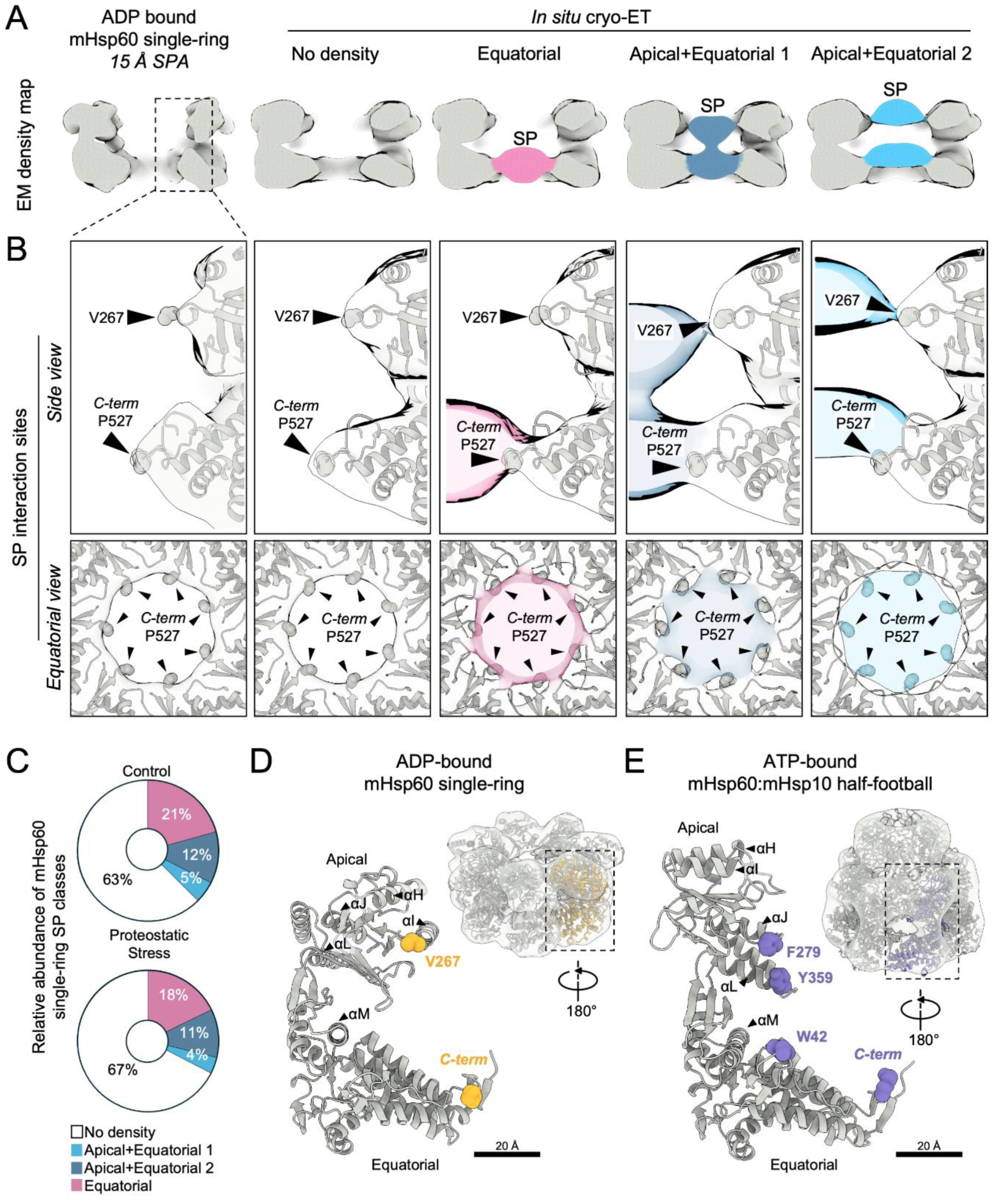
***In situ* mHsp60-substrate protein interactions in mHsp60 single-rings.** (**A**) *In situ* subtomogram averages from focused 3D classification of mHsp60 single-rings (right) compared to a single-particle cryo-EM (SPA) structure of the ADP-bound mHsp60 single-ring low-pass filtered to 15 Å resolution (left). (**B**) Detailed views of SP interactions with mHsp60. The atomic model of the ADP-bound mHsp60 single-ring was docked by rigid body fitting into the subtomogram averaging maps (semitransparent). SPs are colored as in (A) (semitransparent). mHsp60 residues interacting with SPs are shown as spheres and indicated by arrowheads. Interactions are shown in side view (top) and in top view at the level of the equatorial domains (bottom). (**C**) Relative abundance of SP-binding classes identified *in situ*. (**D**) Atomic model of a mHsp60 monomer within the ADP-bound mHsp60 single-ring, with SP-interacting mHsp60 residues highlighted as yellow spheres. The inset shows the atomic model of the ADP-bound single-ring docked in the *in situ* subtomogram averaging map of the mHsp60 single-ring rotated 180° along the vertical axis. (**E**) Atomic model of a mHsp60 monomer within the ATP-bound mHsp60:mHsp10 half-football, with SP-interacting mHsp60 residues highlighted as purple spheres (see Figure 6). The inset shows the atomic model of the ATP-bound mHsp60:mHsp10 half-football docked into the *in situ* subtomogram averaging map of the mHsp60:mHsp10 half-football rotated 180° along the vertical axis.

Next, we investigated the regions of contact between mHsp60 single-rings and their SPs. To that end, we first verified whether SP binding induced any major structural change in the mHsp60 single-ring. However, the “no density” class was comparable to an average of all SP-bound classes pooled (Figure S 6C), and both classes fitted best the ADP-bound single particle cryo-EM structure (Figure S 6E). Thus, the majority of mHsp60 single-rings detected in cells were likely bound to ADP irrespective of whether or not they interacted with SPs. Docking of the atomic model of the ADP-bound mHsp60 single-ring into the *in situ* SP-bound structures of mHsp60 single-rings revealed the major sites of SP interaction (Figure 5B). The “equatorial” class, showed SP contacts with the mHsp60 C-terminal tails, consistent with previous work (Braxton *et al*., 2024). Similar equatorial densities were also observed in the “apical+equatorial” classes, which also displayed an additional contact in the apical domain, corresponding to the loop region (L265-V271) at the end of Helix I (Figure 5D). Comparable contacts were observed also in C1 classes, ruling out symmetry-induced artifacts (Figure S 6D). Therefore, our *in vitro* and *in situ* data suggest that the majority of mHsp60 single-rings in cells are bound to ADP and are able to engage in stable interactions with SPs.

### Substrate protein interactions within mHsp60:mHsp10 complexes

Upon capture of SPs by mHsp60, mHsp10 binding leads to SP encapsulation for subsequent folding (Braxton *et al*., 2024; Gomez-Llorente *et al*., 2020; Nisemblat *et al*., 2015; Weiss *et al*., 2016). Therefore, our mHsp60:mHsp10 complexes detected in cells may have captured some of these interactions. To analyze this possibility, we first pooled all mHsp60:mHsp10 complexes detected in both footballs and half-footballs to obtain an average structure at ∼11 Å nominal resolution (Figure S *7*A, B). This strategy allowed a fair comparison of SP occupancy of football and half-football complexes, given the different abundance of these complexes (Figure 2E). Next, we carried out an analogous classification of mHsp60:mHsp10 complexes as for mHsp60 single-rings, using a mask focused on the central mHsp60:mHsp10 cavity (Figure S *7*C). Again, this resulted in four classes, one of which displayed no SP density, and another three where the SP was visible at the different positions of the chamber (“equatorial”, “apical+equatorial” and “apical”; Figure 6A, Figure S *7*D-F). Similar results were obtained upon independent rounds of classification (Figure S *7*D, E). As discussed above, the class with no visible SP density may represent empty mHsp60:mHsp10, or SPs that are too disordered to be captured by averaging, i.e. SPs in the initial stages of the folding cycle. Whereas we cannot differentiate between these possibilities here, our recent *in situ* cryo-ET work on bacterial GroEL:GroES complexes would be consistent with the latter scenario (Wagner *et al*., 2024). In any case, the fraction of mHsp60:mHsp10 complexes with defined SP densities (Figure 6C) was approximately double than for mHsp60 single-rings (Figure 5C), indicative of a more advanced stage in the SP folding cycle.

**Figure 6:**
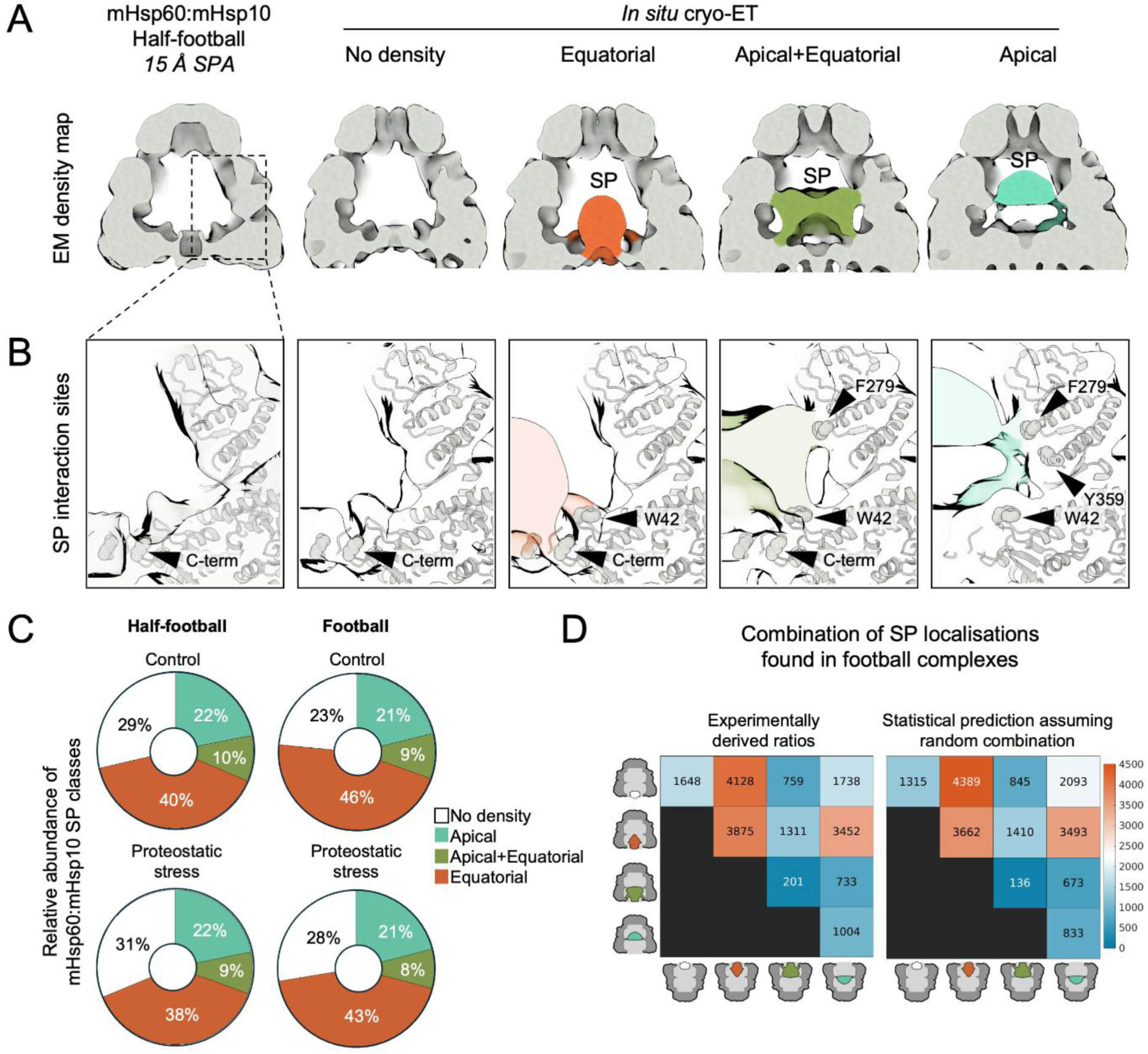
*In situ* mHsp60- substrate protein interactions in mHsp60:mHsp10 complexes. (**A**) *In situ* subtomogram averages from focused 3D classification of SP-bound mHsp60:mHsp10 complexes rings (right) compared to a single-particle cryo-EM (SPA) structure of the ATP-bound mHsp60:mHsp10 half-football low-pass filtered to 15 Å resolution (left). (**B**) Detailed views of SP interactions with mHsp60. The atomic model of the ATP-bound mHsp60:mHsp10 half-football was docked by rigid body fitting in the subtomogram averaging maps (semitransparent). SPs are colored as in (A) (semitransparent). mHsp60 residues interacting with SPs are shown as spheres and indicated by arrowheads. (**C**) Relative abundance of SP-binding classes identified *in situ*. (**D**) Analysis of possible cooperativity between the two halves of mHsp60:mHsp10 football complexes. The experimentally observed distribution of SPs in the two folding cages of footballs (left) is compared to a random distribution (right). The color scale represents the number of particles in each category from a total of the 18,845 mHsp60:mHsp10 football complexes detected in the complete dataset.

Beyond the configuration of the SP, no major structural differences were observed between the classes (Figure 6A, Figure S *7*D, E). Therefore, we docked the atomic models of ATP-bound mHsp60:mHsp10 derived from the single-particle cryo-EM maps into all *in situ* classes to investigate the sites of SP interaction. This revealed that, equatorially, SPs contacted the mHsp60 C-terminal tails, as well as the N-terminal W42 residue (Figure 6B), which have been implicated in SP binding (Braxton *et al*., 2024). SPs contacted the apical domains in a different location (helices J and L; Figure 5E, Figure 6B) compared to mHsp60 single-rings, as helices H and I were no longer available for SP interactions upon mHsp10 binding (Braxton *et al*., 2024). As for mHsp60 single-rings, similar SP interactions were observed in mHsp60:mHsp10 complexes with or without the application of symmetry in the reconstructions (Figure S *7*F).

No major differences in SP binding were found between cells under control and proteostatic stress conditions (Figure 6C), further indicating that mHsp60-SP interactions were not affected by proteostatic stress. SP-bound classes were similarly abundant in mHsp60:mHsp10 footballs and half-footballs, although footballs displayed a slightly higher abundance of classes with ordered SP densities (Figure 6C). These findings suggest that, although mHsp60:mHsp10 half-footballs may be able to carry out the complete folding cycle, they preferentially associate into footballs, especially under proteostatic stress (Figure 2E). Therefore, mHsp60:mHsp10 footballs may be generally more advanced in the folding cycle than half-footballs.

Finally, we investigated possible correlations in SP folding between the two chambers of mHsp60:mHsp10 footballs. To that end, we compared the experimentally observed distribution of SP classes on each of the two halves of any given mHsp60:mHsp10 football to a random distribution. This analysis did not reveal major differences (Figure 6D), suggesting that the two halves of mHsp60:mHsp10 footballs operate independently from each other. Given that neither mHsp60 double rings nor mHsp60:mHsp10 bullets were observed in cells, our data imply that within mitochondria mHsp60:mHsp10 footballs are mainly formed by equatorial association of two ATP-bound half-footballs. ATP hydrolysis in one of the halves may disassemble the football to initiate a new folding cycle (Figure 4E, Figure *7*B).

**Figure 7:**
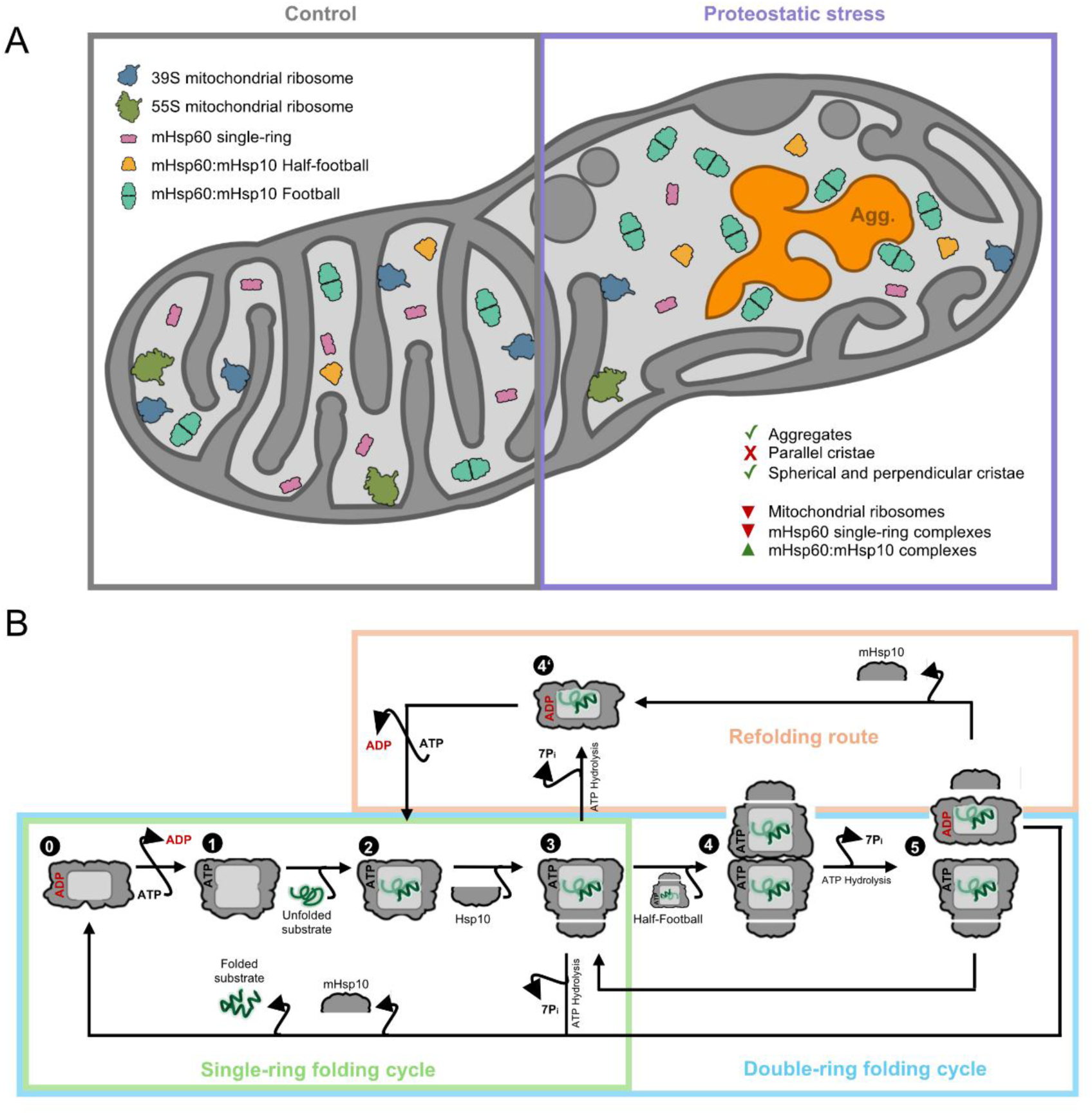
Structural models for mitochondrial remodeling upon proteostatic stress and the mHsp60 functional cycle. (**A**) Summary of the major structural rearrangements observed in mitochondria upon proteostatic stress, including changes in mitochondrial morphology and remodeling of mitochondrial ribosome and mHsp60 complexes. (**B**) Proposed model for the native mHsp60:mHsp10 SP folding cycle in mitochondria, encompassing both single-ring and double-ring folding pathways, as well as a potential refolding route for SPs requiring multiple rounds assisted folding.

## Discussion

Our cryo-ET analyses reveal *in situ* structural consequences of mitochondrial proteostatic stress (Figure 7). Our data suggest that protein aggregates in the mitochondrial matrix alter cristae organization, first locally and eventually throughout the entire organelle (Figure *7*A). Importantly, both mHsp90 inhibition (this study) and the import of aggregation-prone proteins into mitochondria (Holthusen *et al*., 2025) led to a similar phenomenon, suggesting that cristae disruption is a general consequence of protein aggregation in the matrix. Although little is known about the effects of proteostatic stress on cristae morphogenetic factors, alterations in cristae shape may impact the stability of respiratory complexes (Cogliati *et al*, 2013), favoring their sequestration into aggregates. This suggests a vicious cycle, in which increased aggregation of respiratory complexes further disrupts cristae organization, leading to further destabilization of respiratory complexes. This phenomenon may underlie the reported loss of mitochondrial respiratory capacity under proteostatic stress (Holthusen *et al*., 2025; Uoselis *et al*., 2023).

Previous work showed that G-TPP-induced proteostatic stress activates the UPR^mt^ leading to reduced mitochondrial translation (Fiesel *et al*., 2017; Kang *et al*., 2009; Munch & Harper, 2016; Uoselis *et al*., 2023). Our cryo-ET data rationalize these findings by showing a strong reduction in mitochondrial ribosome complexes under proteostatic stress (Figure 2B). Furthermore, our analyses hint to an active role of protein aggregation in mitochondrial ribosome depletion, as significantly lower numbers of fully assembled 55S ribosomes were found in the vicinity of aggregates (Figure 2H, Figure *7*A). These data are consistent with recent evidence that mitochondrial aggregates sequester mitochondrial ribosome components, especially assembly factors and proteins of the small ribosomal subunit (Bertgen *et al*, 2024; Holthusen *et al*., 2025). Thus, mitochondrial ribosome assembly appears to be particularly sensitive to proteostatic stress within the matrix.

The mitochondrial chaperonin mHsp60:mHsp10 assists the folding of about half of the matrix proteome, is crucial for the assembly of mitochondrial ribosomes, and is a key UPR^mt^ effector (Bie *et al*., 2020; Holthusen *et al*., 2025; Martinus *et al*., 1996; Munch & Harper, 2016; Yoneda *et al*., 2004; Zhao *et al*., 2002). Building on our combination of *in situ* and *in vitro* structural analyses, we propose a model for the mHsp60 functional cycle within cells and its modulation by misfolding stress (Figure *7*B). We find that under basal conditions mHsp60 single-rings predominate (Figure 2E, F), in most cases without a defined density for SPs (Figure 5C) and likely bound to ADP (Figure 3B, Figure S 4C, D, Figure S 6E; Figure *7*B, stage 0). Consistently, the cryo-EM structures suggest that ADP-bound mHsp60 single-rings display lower affinity for SPs than ATP-bound mHsp60 single-rings, given the positioning of SP-interacting helices H and I away from the central cavity of the ring, as well as the reduced dynamics of the apical domains in the ADP-bound form (Figure 3, Figure 4, Figure S 5B) (Braxton *et al*., 2024; Tascon *et al*., 2025). We suggest that the abundance of ADP-bound mHsp60 single-rings in cells may reflect i) the relatively long residency time of ADP (Mas *et al*, 2018), and ii) the rapid binding dynamics of SPs and the mHsp10 protein cofactor following ADP exchange by ATP (Figure *7*B, stages 1, 2). In this framework, ATP-bound mHsp60 single-rings in cells are likely too transient to be detected by our approach, and the next stable (and thus detectable) assembly is the ATP-bound mHsp60:mHsp10 half-football encapsulating a SP (Figure *7*B, stage 3). Most mHsp60:mHsp10 half-football particles displayed ordered SP densities within their central cages (Figure 6), indicating that these SPs were at least partly folded. This implies that mHsp60:mHsp10 half-footballs are competent for SP folding in mitochondria, consistent with early reports (Nielsen & Cowan, 1998) and our previous findings that obligated half-football mHsp60:mHsp10 variants can complement GroEL:GroES-deficient *E. coli* strains (Gomez-Llorente *et al*., 2020). After an attempted folding event, ATP hydrolysis by the mHsp60:mHsp10 half-footballs may trigger mHsp10 release (Nielsen & Cowan, 1998), and two outcomes may follow: if folding is successful, the folded SP would be released along with mHsp10, regenerating an ADP-bound mHsp60 single-ring ready to reinitiate the cycle (Figure *7*B, stage 0). This pathway represents the single-ring folding cycle (Figure *7*B, green rectangle).

However, it is known that SPs often require multiple rounds of chaperonin engagement to achieve full folding (Lin *et al*, 2013; Weissman *et al*, 1994). Releasing partially misfolded SPs after each cycle would be inefficient, as it would necessitate repeated SP capture events and could promote deleterious aggregation (Hayer-Hartl *et al*., 2016). We therefore propose that if SPs remain incompletely folded upon ATP hydrolysis and mHsp10 release, a fraction of them may remain bound to the ADP-bound mHsp60 single-ring, awaiting nucleotide exchange and mHsp10 rebinding for another folding cycle (Figure *7*B, stage 4’). In this model, the ∼35% of mHsp60 single-rings observed bound to an ordered SP (Figure 5C) would primarily represent SPs undergoing refolding (Figure *7*B, pink rectangle). The visible SP densities suggest that SPs are already partially ordered, and may thus have experience previous folding rounds. Thus, these densities are less likely to arise from newly captured SPs, which would generally be more disordered and preferentially bind to empty ATP-bound mHsp60 single-rings. This “refolding route” may be favored for SPs with intrinsically slow folding kinetics.

Under proteostatic stress, most mHsp60:mHsp10 complexes detected in cells corresponded to footballs with two folding cages (Figure 2E, F). Therefore, although mHsp60:mHsp10 half-footballs are likely folding-competent, they appear to preferentially associate equatorially into footballs (Figure *7*B, stage 4). This is consistent with the ability of ATP to flatten the mHsp60 equatorial interface (Figure 4E) and promote inter-ring dimerization (Levy-Rimler *et al*., 2001; Tascon *et al*., 2025). Ordered SP densities were slightly more abundant within mHsp60:mHsp10 footballs than within half-footballs (Figure 6C), in agreement with footballs resulting from the association of two half-footballs in which folding had already started. The enrichment of football complexes within the immediate vicinity of mitochondrial aggregates (Figure 2H, Figure *7*A) suggests that the abundance of misfolded SPs promotes football assembly. Our analyses of the two halves of mHsp60:mHsp10 footballs indicated that each folding cage operates independently without detectable cooperativity, unlike GroEL:GroES (Hayer-Hartl *et al*., 2016; Wagner *et al*., 2024).

Because mHsp60:mHsp10 footballs form through association of ATP-bound half-footballs at different stages of the folding cycle, ATP hydrolysis is likely to occur first in one half. This would convert ATP to ADP, destabilizing the equatorial interface and triggering the split of the football (Figure *7*B, stage 5). This event marks the end of the double-ring folding cycle (Figure *7*B, blue rectangle). The resulting ADP-bound half-football would then release mHsp10 as discussed above, and either discharge its folded SP to restart the cycle (Figure *7*B, stage 0), or retain a partially-folded SP for subsequent refolding (Figure *7*B, stage 4’). The other half-football remains ATP bound (Figure *7*B, stage 3), and may thus continue the folding cycle either as an independent half-football or by associating into a new football complex.

An alternative pathway for football assembly could involve preformed mHsp60 double rings binding two mHsp10 complexes. Given that simultaneous binding of both mHsp10 lids is unlikely, this route would probably proceed via a “bullet” intermediate, where one ring is capped by mHsp10 and the other is not. Although we resolved the structures of such double-ring and bullet complexes *in vitro* (Figure S 5), neither species was detected in cells despite a targeted search. Thus, while transient formation of mHsp60 double-rings and mHsp60:mHsp10 bullets cannot be excluded, our data indicate that they are unlikely to represent a major pathway for mHsp60:mHsp10 football formation in cells. In contrast, we have recently shown that GroEL:GroES bullets are the most abundant conformation within *E. coli* cells (Wagner *et al*., 2024). Therefore, despite their high sequence and structural homology, bacterial GroEL:GroES and human mHsp60:mHsp10 populate distinct assembly landscapes throughout their SP folding cycles.

Collectively, our work sheds light into the functional mechanisms of key mitochondrial machineries for protein biogenesis and their remodeling in response to proteostatic stress.

### Limitations of this study

This study focused on two key machineries for mitochondrial protein biogenesis: the mitochondrial ribosome and the mHsp60 chaperonin system. However, mitochondrial proteostatic responses also engage various other protein quality control factors, including mHsp70, mHsp90 and proteases whose structural transitions upon stress are yet to be elucidated. The nature of 3D classification gives us confidence that mHsp60 single-rings, mHsp60:mHsp10 half-footballs and mHsp60:mHsp10 footballs are the most populated mHsp60 assemblies in our cells, but we cannot exclude small populations of other mHsp60 assemblies. Similarly, although we detected only ADP-bound mHsp60 single-rings in cells, other nucleotide states likely exist, albeit transiently. Furthermore, given that mHsp60:mHsp10 half-footballs are likely competent for folding SPs, it remains to be determined why football formation is favored, and whether a football is more folding-competent than two separate half- footballs. Lastly, our data complement recent cryo-ET analyses of mitochondrial alterations in the context of mitochondrial depolarization that also induce PINK1 activation (Rose *et al*, 2025). Future studies directed at visualizing PINK1 accumulation at the TOM complex under mitochondrial stress may aid in understanding how these alterations signal mechanistically to PINK1 activation.

## Resource availability

### Lead contact

Further information and requests for resources and reagents should be directed to and will be fulfilled by the lead contact, Rubén Fernández-Busnadiego.

### Materials availability

HeLa PINK1-GFP knock-in cells are available upon request. No other unique reagents were generated in this study.

### Data and code availability

Structural data generated in this study have been deposited at EMDB (density maps) and PDB (atomic models) and is available as follows: *in situ* subtomogram averaging density maps of the 39S mitochondrial ribosome (EMDB 55203), 55S mitochondrial ribosome (EMDB 55204), mHsp60 single-ring (EMDB 55205), mHsp60 half-football (EMDB 55206) and mHsp60 football (EMDB 55207); single-particle cryo-EM density maps and atomic models of the ATP-bound mHsp60:mHsp10 football complex (D7 symmetry; EMDB 54898, PDB 9SHG), ATP-bound mHsp60:mHsp10 half-football complex (C7 symmetry; EMDB 54900, PDB 9SHI), ATP-bound mHsp60:mHsp10 bullet complex (C7 symmetry; EMDB 54899, PDB 9SHH), ATP-bound mHsp60 double-ring complex (C7 symmetry; EMDB 54901, PDB 9SHJ), ATP-bound mHsp60 single-ring complex (C7 symmetry; EMDB 54903, PDB 9SHL) and ADP-bound mHsp60 single-ring complex (C7 symmetry; EMDB 54902, PDB 9SHK).

Code generated in this study (PROCMAN and MuRePP scripts) is available at https://gitlab.gwdg.de/fernandez-busnadiego-lab/mitochondrial-proteostasis-cryo-et-analysis

## Acknowledgments

We are grateful to P. Kakade for the generation of HeLa PINK1-GFP knock-in cells, T. Cheng for assistance in cryo-ET experiments, C. Weiss for assistance with mHsp60 and mHsp10 purification, J. Wagner and A. Petrovic for advice on cryo-ET data processing, as well as E. Sakata, F.U. Hartl, W. Harper, I. Tascón, A. G. Berruezo, J. Vilchez-Garcia, C. Weiss andJ.A. Hirsch for fruitful discussions.

This work was funded by the Deutsche Forschungsgemeinschaft (DFG, German Research Foundation) under Germany’s Excellence Strategy (EXC 2067/1-390729940 to R.F.-B.), Wellcome Trust through a Senior Clinical Fellowship (210753/Z/18/Z) to M.M.K.M., the Michael J. Fox Foundation for Parkinson’s Research (MJFF) through grant MJFF-010458 to M.M.K.M. and R.F.-B., and the joint efforts of the MJFF and the Aligning Science Across Parkinson’s (ASAP) initiative. MJFF administers grants ASAP-000463 (to M.M.K.M.) and ASAP-000282 (to R.F.-B.) on behalf of ASAP and itself. Furthermore, A.A. acknowledges funding from the Israel Science Foundation (1723/23) and the Recanati Foundation from Tel Aviv University. M.M.K.M acknowledges funding from the Medical Research Council and the UK Dementia Research Institute and Parkinson’s UK. I.U.-B. acknowledges funding from the Spanish Ministry of Science, Innovation and Universities (MICIU), the Spanish Research Agency (AEI) and the European Regional Development Fund (MICIU / AEI / 10.13039/501100011033 / FEDER, UE) through grant PID2022-143177NB-I00. J.P.L.-A. acknowledges financial support from a PTA contract granted by the MICIU / AEI.

Cryo-ET instrumentation at the University of Göttingen was jointly funded by the DFG Major Research Instrumentation program (448415290) and the Ministry of Science and Culture of the State of Lower Saxony. Single-particle cryo-EM data collection was performed at the Basque Resource for Electron Microscopy located at Instituto Biofisika (UPV/EHU, CSIC), supported by the Department of Science, Universities and Innovation and the Innovation Fund of the Basque Government, by Fundación Biofísica Bizkaia and with additional support from MICIU (Recovery, Transformation and Resilience Plan) and the Basque Government “Biotechnology Complementary Plan Applied to Health” with funding from European Union NextGenerationEU (PRTR-C17.I1; PRTR-C17.I01.P01.S13; AAAA_ACG_AY_2539/22_05).

## Author contributions

K.E. carried out cell biological and cryo-ET experiments, including sample preparation, data collection and analysis. J.P.L.-A. carried out structural determination by single-particle cryo-EM. O.A. carried out biochemical experiments. A.A. provided expertise in purification of mHsp60 and mHsp10. M.M.K.M. supervised biochemical work. I.U.-B. supervised single-particle cryo-EM work. R.F.-B. supervised cryo-ET work. A.A., M.M.K.M., I.U.-B. and R.F.-B. planned research. K.E., I.U.-B. and R.F.-B. wrote the manuscript with contributions from all other authors.

## STAR Methods

### Protein expression and purification

mHsp60 is translated in the cytosol as a 573-amino acid polypeptide containing a 26- amino acid long mitochondrial matrix targeting signal, which upon cleavage yields a mature 547 amino acid long mitochondrial protein beginning with the sequence AKDVKFG. For recombinant mHsp60 production, the WT human mHsp60 gene (*HSPD1*), lacking the N-terminal mitochondrial targeting sequence, was expressed in *E. coli* as a construct bearing an N-terminal HisTag followed by a TEV protease cleavage site. The protein was purified by sequential Ni-affinity and anion-exchange chromatography, as previously described (Gomez-Llorente *et al*., 2020; Weiss *et al*, 2024). After TEV cleavage, the resulting mHsp60 used for cryo-EM retained an additional Gly-Ser at the N- terminus that does not affect the final structure or function of the reconstituted protein (Gomez-Llorente *et al*., 2020). For consistency with recent cryo-EM studies (Braxton *et al*., 2024; Klebl *et al*., 2021; Wang & Chen, 2021), amino acid numbering on our structures refers to the mature WT mitochondrial mHsp60.

Monomeric mHsp60 was concentrated and incubated at 30 °C for 2 h in the presence of 4 mM ATP, 20 mM KCl, and 20 mM magnesium acetate to promote oligomer assembly. The sample was then loaded onto a Superdex 200 gel-filtration column equilibrated in 50 mM Tris-HCl pH 7.7, 300 mM NaCl, and 10 mM MgCl2. Fractions containing active oligomeric mHsp60 were pooled, concentrated, and flash-frozen in liquid nitrogen for storage. WT human mHsp10 (*HSPE1*) was expressed in *E. coli* without a HisTag and purified by sequential anion-exchange and cation-exchange chromatography followed by gel filtration, as previously reported (Gomez-Llorente *et al*., 2020). Purified mHsp10 in 50 mM Tris-HCl pH 7.7 and 100 mM NaCl was concentrated and flash-frozen in liquid nitrogen for storage.

### Sample preparation and data collection for single-particle cryo-EM

To capture mHsp60:mHsp10 assembly intermediates as a function of ATP binding and hydrolysis, mHsp60 and mHsp10 were mixed at a final concentration of 10 μM and 8 μM, respectively (molar ratio 1:0.8), in reaction buffer containing 20 mM Tris-HCl pH 7.7, 20 mM KCl, 10 mM MgCl₂ and 2 mM ATP. After mixing, samples were incubated for defined time intervals ranging from 30 s to 30 min. At each time point, a 4 μl aliquot of the reaction mixture was applied to glow-discharged QuantiFoil R 2/1 300 mesh grids. Grids were blotted from the front side for 2.4-2.7 s under 95% relative humidity and vitrified in liquid ethane using a Leica GP2 cryo-plunger (Leica Microsystems).

Specimens were imaged on a 300 kV Krios G4 (Thermo Fischer Scientific) cryo-electron microscope (cryo-TEM) equipped with a BioContinuum energy filter and a K3 direct detector camera (Gatan) operating in counting mode at a calibrated 0.8238 Å per pixel. Employing a 0.8-1.6 μm defocus range, six movies were recorded per hole, each with a total accumulated dose of 50 e^−^/Å^2^ over 50 frames. Movies were recorded automatically using EPU 2 (Thermo Fischer Scientific) with aberration-free image shift and fringe-free imaging.

### Cryo-EM data processing and 3D reconstruction

Initial processing for datasets collected at different time points followed an analogous general strategy (Figure S 5). Frame alignment and contrast transfer function (CTF) estimation were performed in cryoSPARC Live (Punjani *et al*, 2017). Micrographs were evaluated based on CTF fits and total motion, and the best movies were selected for further processing. For each dataset, automatic particle picking was carried out in cryoSPARC using templates generated from an initial blob-based picker. Several rounds of reference-free 2D classification were used to remove poorly aligning particles.

*Ab initio* reconstruction was performed to generate initial 3D models. Particles contributing to mHsp60 single-ring classes in the *ab initio* reconstructions were further subjected to additional *ab initio* reconstructions to obtain distinct initial mHsp60 single-ring models. Heterogeneous refinement was then performed to sort particles among the different conformational states. For each of the resulting classes, homogeneous refinements were carried out separately.

After processing the datasets individually, particles assigned to the same structural state across different time points were combined. The merged particle stacks were further cleaned by additional rounds of 2D classification and heterogeneous refinement to remove outliers. Final refinements were performed in RELION 3.0 with per-particle CTF refinement enabled (Zivanov *et al*, 2020). The local resolution of the reconstructions was estimated in cryoSPARC.

### Model building

For model building of the ATP-bound mHsp60:mHsp10 football, mHsp60:mHsp10 half-football, and mHsp60 double-ring complexes, we used as starting models our previously deposited structures with PDB IDs 9ES0, 9ES1, and 9ES2, respectively (Tascon *et al*., 2025). For the ATP-bound mHsp60 single-ring complex, we generated an initial model by extracting one ring from the mHsp60 double-ring ATP structure (PDB 9ES2). The mHsp60:mHsp10 bullet-shaped complex was assembled by combining a mHsp60:mHsp10 half-football models (PDB 9ES1) with one ring of the ATP-bound mHsp60 double-ring structure. For the ADP-bound mHsp60 single-ring complex, we used the mHsp60:mHsp10 half-football model as the starting template.

Rigid-body fitting of the initial models into the corresponding cryo-EM density maps was performed using UCSF ChimeraX (Goddard *et al*., 2018). Subsequent rounds of real-space refinement were carried out in Phenix (Adams *et al*, 2010) with secondary structure and geometry restraints. Manual model correction and rebuilding were performed in Coot (Emsley & Cowtan, 2004) to improve the fit and stereochemistry prior to final refinement. RMSD calculations were performed using the Matchmaker tool within ChimeraX (Goddard *et al*., 2018).

In the ATP-bound mHsp60 single-rings, intermediate and equatorial domains were well resolved, while apical densities were weak, in line with previous results (Braxton *et al*., 2024; Tascon *et al*., 2025). Gaussian filtering of the cryo-EM map in Chimera (Pettersen *et al*, 2004) allowed visualization of the apical region density (Figure S 5B). Thus, the structure of the equatorial and intermediate domains of the ATP-bound mHsp60 single-ring was built *de novo* based on the high-resolution density features of the cryo-EM map in this region, whereas for the apical region the analogous region of an apo mHsp60 single-ring structure (PDB 9ES3) (Tascon *et al*., 2025) was docked and refined.

### Generation of HeLa PINK1-GFP knock-in cell line using CRISPR/Cas9

A HeLa cell line expressing PINK1 fused to GFP at the C-terminus from the endogenous locus was generated via CRISPR/Cas9. A double-strand break was introduced in exon 1 of the PINK1 gene using two sgRNAs delivered on Cas9-nickase plasmids (DU64062 and DU64072), along with a donor plasmid (DU64119) encoding full-length PINK1- GFP, a stop codon, and a polyadenylation sequence. The donor was flanked by ∼500 bp homology arms. C-terminal tagging was chosen to preserve the N-terminal mitochondrial targeting sequence.

HeLa cells (∼60% confluency) were transfected in 10 cm dishes using polyethylenimine (PEI) in antibiotic-free Dulbecco’s modified Eagle’s medium (DMEM) with 10% fetal bovine serum (FBS) and 2 mM L-glutamine. A total of 1 µg each of the CRISPR plasmids and 3 µg of the donor plasmid were mixed in 1 mL of Opti-MEM with 20 µL PEI, incubated for 30 min at room temperature, and added to the cells. After 24 h, puromycin (2 µg/mL) was applied for 48 h. A second transfection was performed without selection, followed by a 10-day culture period to eliminate residual donor expression.

To boost PINK1 accumulation and enhance GFP detection, cells were treated with 5 µM antimycin A and 0.63 µM oligomycin for 18 h, then washed and cultured for another 24 h. GFP-positive cells were sorted by FACS (530/540 nm filter) using a low GFP threshold to exclude clones with off-target integration or defective PINK1 turnover. Sorted cells were plated in 96-well plates coated with 0.1% gelatine. Single colonies were expanded (5-6 weeks) and screened by immunoblotting with anti-PINK1 and anti-GFP antibodies. A detailed protocol for these procedures is available at dx.doi.org/10.17504/protocols.io.dm6gpm38pgzp/v1.

### Whole-cell lysate preparation

For biochemical experiments, HeLa GFP-PINK1 cells were cultured in DMEM supplemented with 10% FBS, 1% penicillin/streptomycin, and 1% L-glutamine, and used for experiments at 90% confluency. Cells were treated with 10 µM G-TPP, and PINK1 activation was assessed at 1, 2, 3, 4, 5, and 6 h. DMSO was used as control. HeLa GFP-PINK1 cells were sonicated in lysis buffer containing 50 mM Tris·HCl (pH 7.5), 1 mM EDTA, 1 mM EGTA, 1% Triton (w/v), 0.1% SDS (w/v), 1 mM sodium orthovanadate, 10 mM sodium glycerophosphate, 50 mM sodium fluoride, 10 mM sodium pyrophosphate, 0.25 M sucrose, 0.1 mM phenylmethylsulfonyl fluoride, protease inhibitor cocktail (Roche), phosphatase inhibitor cocktails 2 (sigma) and phosStop (Roche), and 200 mM chloroacetamide. After sonication, lysates were incubated on ice for 30 min. Samples were centrifuged at 17,000 g for 30 min at 4 °C using an Eppendorf 5417R centrifuge. The supernatants were collected, and protein concentration was determined using BCA (Pierce). A detailed protocol for these procedures is available at dx.doi.org/10.17504/protocols.io.bswanfae.

### Ubiquitin capture using Halo-m-DSK

Halo-tagged m-DSK, a yeast Dsk1-derived ubiquitin-binding protein, was immobilized by incubating with 200 μL HaloLink resin (Promega) in binding buffer (50 mM Tris-HCl [pH 7.5], 150 mM NaCl, 0.05% NP-40) overnight at 4 °C. Whole-cell lysates (1 mg) were then incubated with 20 μL of Halo-m-DSK-bound resin overnight at 4 °C. Beads were washed three times with lysis buffer containing 0.25 M NaCl, and bound proteins were eluted by resuspending in 20 μL of 2× lithium dodecyl sulfate (LDS) sample buffer. The eluates were incubated at 37 °C for 15 min with shaking (2000 rpm), followed by the addition of 2.5% 2- mercaptoethanol. This method was adapted from a previously described m-DSK ubiquitin capture protocol (Wilson *et al*, 2012).

### GFP-PINK1 immunoprecipitation and immunoblotting

Whole-cell lysates (1 mg) were incubated overnight at 4 °C with 10 μg of GFP beads (DU61915Agarose-MRC Reagent & Services). Beads were washed three times with lysis buffer containing 250 mM NaCl, and bound proteins were eluted in 10 μL of 2× LDS sample buffer. Eluates were heated at 70 °C for 10 min, followed by the addition of 2.5% 2- mercaptoethanol. This protocol was adapted from dx.doi.org/10.17504/protocols.io.eq2ly7kxqlx9/v1.

Samples were resolved by SDS–PAGE using 4–12% Bis-Tris gels and transferred to 0.45 μm polyvinylidene difluoride membranes (Protran, Immobilon-P). Membranes were blocked for 1 h at room temperature in 5% skimmed milk in TBS-T (50 mM Tris-HCl, 150 mM NaCl, 0.1% Tween-20, pH 7.5), and incubated overnight at 4 °C with the indicated primary antibodies. Detection was performed using HRP-conjugated secondary antibodies and enhanced chemiluminescence. A detailed protocol for PINK1-Parkin signalling immunoblotting is available at dx.doi.org/10.17504/protocols.io.bswanfae.

### Fluorescence microscopy

Cells were seeded onto poly-L-lysine–coated glass coverslips placed in 24-well plates and allowed to adhere overnight. The following day, cells were either left untreated or treated with 10 µM G-TPP for the indicated durations. 30 min prior to the end of the incubation period, 300 nM MitoSpy Red CMXRos (BioLegend) was added to stain mitochondria. Cells were then washed with PBS and fixed with 4 % paraformaldehyde for 15 min at room temperature. After three washes in PBS to remove residual fixative, cells were permeabilized with 0.1 % Triton X-100 and incubated for 5 min with Hoechst 33342 to stain nuclear DNA. Coverslips were extensively washed with PBS and mounted on glass slides using a compatible mounting medium.

Fluorescence imaging was performed on a DMi8 microscope (Leica) equipped with an HC PL APO 63×/1.40 oil immersion objective and a Leica K5-14401415 detector. Image acquisition was carried out as 2×2 binned Z-stacks comprising 20 steps and three channels corresponding to the respective signals (MitoSpy Red CMXRos in the TXT channel, PINK1- GFP in the GFP channel, and DAPI in the DAPI channel). Further processing, including histogram adjustment and denoising with THUNDER, was performed using the LAS X software platform (Leica).

### Cell viability assay

Cell viability was assessed using the CellTiter-Glo 2.0 assay (Promega) according to the manufacturer’s instructions. In brief, cells were seeded into 96-well flat-bottom white opaque plates and allowed to adhere overnight. Following treatment with 10 µM G-TPP for the indicated durations, CellTiter-Glo reagent was added directly to the wells and incubated for 10 min at room temperature. Luminescence was then measured using a plate reader (Tecan). Experiments were performed in triplicate.

### Cryo-ET sample preparation

Electron microscopy 200-mesh R2/2 gold Ultrafoil grids (Quantifoil) were glow-discharged and immersed in cell culture medium. Cells were plated directly onto the grids with a target confluency of ∼30–50 % and allowed to adhere overnight. The following day, cells were either left untreated or treated with 10 µM G-TPP for 4 h prior to vitrification using a Vitrobot Mark IV (Thermo Fisher Scientific).

Immediately before freezing, grids were incubated in 10 % glycerol for ∼30 s. Blotting was performed from the back under the following conditions: 37 °C chamber temperature, blot force 0, blot time 5 s, drain time 1 s, without humidification. Grids were then plunge-frozen into a liquid ethane/propane mixture and stored in liquid nitrogen in cryo-EM storage boxes until further use.

### Cell thinning by focused ion beam–scanning electron microscopy (FIB/SEM)

Grids were clipped into autogrids featuring FIB-compatible cutouts and mounted onto the cryo-FIB/SEM transfer shuttle. Lamella preparation was performed using an Aquilos 2 cryo-FIB/SEM dual-beam microscope (Thermo Fisher Scientific). Prior to thinning, samples were sequentially coated with protective platinum layers: an initial sputter-coated layer of inorganic platinum, followed by an organometallic platinum deposition, and a final sputter-coated layer of inorganic platinum. Full-grid SEM montages were acquired to identify suitable cells for lamella preparation.

Automated rough milling down to a lamella thickness of ∼400 nm was performed using the AutoTEM 5 software (Thermo Fisher Scientific) in a stepwise fashion with decreasing ion beam currents (1 nA to 300 pA) at a milling angle of 10°. Final polishing was carried out manually using currents ranging from 100 to 30 pA to achieve a final lamella thickness of <250 nm. After thinning, grids were retracted from the microscope and stored in cryo-EM boxes under liquid nitrogen until further use.

### Cryo-ET data collection

Cryo-electron tomography data were acquired on a Titan Krios G4 transmission electron microscope (Thermo Fisher Scientific) operated at 300 kV. The instrument was equipped with an XFEG electron source, a Selectris post-column energy filter, and a Falcon 4i direct electron detector. Low-magnification full-grid montages were acquired at 135× nominal magnification to locate lamellae. For each lamella, overview montages were recorded at 6,500× magnification (18.89 Å/pixel) to identify mitochondria based on their characteristic morphology. Tomographic tilt series were acquired using SerialEM (Mastronarde, 2005) with the PACEtomo implementation (Eisenstein *et al*, 2023) at a nominal magnification of 53,000× (2.3124 Å/pixel). Tilt series were collected using a dose-symmetric tilt-scheme (Hagen *et al*, 2017), starting from a pre-tilt of 9° and proceeding in 3° increments over a range of ±54°. The dose per micrograph was set to ∼4 e⁻/Å², yielding a cumulative electron dose of <150 e⁻/Å² per tilt-series. Defocus was cycled between −3 and −5 µm in 0.25 µm steps. An energy filter slit of 20 eV width was applied during acquisition to improve contrast.

### Cryo-ET tilt-series reconstruction

Pre-processing was performed in a semi-automated manner using PROCMAN, a collection of MATLAB scripts (https://gitlab.gwdg.de/fernandez-busnadiego-lab/mitochondrial-proteostasis-cryo-et-analysis) partially based on the TOMOMAN suite (Khavnekar *et al*, 2024). In brief, dose-fractionated movies were first motion-corrected and aligned using MotionCor2 (Zheng *et al*, 2017). Frames exhibiting blurring or acquisition artifacts (e.g. dark frames) were manually excluded following visual inspection. Curated tilt series were then aligned by patch tracking in IMOD (Mastronarde & Held, 2017), and tomographic volumes were reconstructed using weighted back-projection with binning 4, resulting in a final pixel size of 9.25 Å.

For subsequent integration into the M/WARP pipeline (Tegunov *et al*, 2021), motion-corrected frame stacks and corresponding IMOD alignment models were imported into WARP for CTF estimation.

### Tomogram segmentation and membrane density analysis

To enhance contrast and improve visibility of cellular structures, reconstructed tomograms were processed using CTF deconvolution implemented in IsoNet (Liu *et al*, 2022). Initial membrane segmentations were generated using MemBrain using the pre-trained model as described (Lamm *et al*, 2022). Amorphous aggregates were segmented using the deep learning framework in Dragonfly. For model training, tomograms were prepared following the protocol described by (Heebner *et al*, 2022). In brief, 15 representative tomograms containing aggregates were manually annotated to train a 2.5D U-Net model, which was subsequently used for prediction to generate initial segmentations. Outputs from both MemBrain and Dragonfly were manually reviewed and refined using Amira (Thermo Fisher Scientific). Segmentations of the full mitochondrial volume were generated manually in Amira.

For the analysis of volumetric membrane density (Figure 1C, D), the number of aggregate-associated voxels was quantified relative to the segmented mitochondrial volume. To classify tomograms into high-aggregation and low-aggregation groups, k-means clustering was performed in MATLAB (Mathworks) using both the relative proportion and absolute volume of aggregate voxels (100,000 replicates). For distance-dependent analysis, the Euclidean distance between each annotated membrane voxel and its nearest aggregate voxel was computed. Voxels within defined distance bins were pooled across tomograms, and the mean membrane density per distance range and tomogram was determined.

### Particle localization and sub-tomogram averaging

To generate data-derived reference maps for template matching (TM) and validate the detectability of specific protein complexes, an initial screen was performed on a subset of 15 control and 15 G-TPP-treated tomograms. As a reference for the mitochondrial ribosome, a density map of the human 55S mitochondrial ribosome (EMDB 3784)(Englmeier *et al*, 2017) was used, after manually removing the membrane signal using UCSF Chimera (Pettersen *et al*., 2004) and low-pass filtering the map to 60 Å. For mHsp60 complexes, a map of the human mHsp60:mHsp1010 football complex (EMDB 9195)(Gomez-Llorente *et al*., 2020) served as a basis. Specific substructures (mHsp60 single-rings, mHsp60 double rings as well as mHsp60:mHsp10 half-footballs, bullets, and footballs) were obtained by targeted masking. All maps were low-pass filtered to 40 Å for TM.

TM was performed in DYNAMO (Castano-Diez *et al*, 2012) with an angular sampling of 15° on bin 4 tomograms for mHsp60 complexes and on bin 8 tomograms for the mitochondrial ribosome. To reduce false positives, cross-correlation (CC) maps were masked using the mitochondrial volume segmentation prior to peak extraction. The top-scoring 600 CC peaks (for mHsp60:mHsp10 footballs) or 500 (for all other species) per tomogram were selected. Sub-volumes were reconstructed in WARP with a pixel size of 9.25 Å and subjected to 2–3 rounds of 3D classification in RELION 4.0 (Zivanov *et al*, 2022). If the classification yielded structurally consistent classes (mHsp60 single-rings, mHsp60:mHsp10 half-footballs and mHsp60:mHsp10 footballs), a 3D refinement was performed to generate a data-derived reference for a second round of TM. Targets that did not yield reliable reconstructions (mHsp60 double rings and mHsp60:mHsp10 bullets) were excluded from further analysis.

In the second round of TM, the same parameters were used as for the initial TM round, but the full dataset of 242 tomograms was searched using the data-derived templates. Based on particle yields from the first TM round, the following numbers of top-scoring coordinates were extracted per tomogram: 1000 (mHsp60:mHsp10 football), 120 (mitochondrial ribosome), 500 (mHsp60:mHsp10 half-football), and 350 (mHsp60 single-ring). Sub-volumes were again reconstructed in WARP (9.25 Å/pixel) and subjected to iterative 3D classification in RELION 4.0. This process was then repeated at higher resolution (4.6248 Å/ pixel). Given the high degree of crowding of the mitochondrial matrix, a second round of particle extraction was performed using the same CC maps, but after masking out all previously verified particle localizations to avoid double picking. The newly extracted sub-volumes were processed as before and pooled with the original sets after distance-based cleaning.

To further improve picking accuracy, a deep-learning-based method was applied using DeepFinder (Moebel *et al*, 2021). For each species, coordinates and STA-derived density maps were used to train a model based on shape-based target recognition, with 95% of the particles used for training and 5% for validation. The model was trained for 70 epochs with 200 training and 20 validation steps per epoch. Coordinates were extracted from clusters with a minimum size of 30 voxels and subjected to the same classification pipeline using WARP and RELION 4.0. DeepFinder-derived particles were then combined with the TM-derived sets, followed by distance-based cleaning.

To prevent early-stage misidentification of particle species, all coordinates from mHsp60 complexes were merged into a single set and subjected to a final round of 3D classification in RELION 4.0. The same procedure was applied to all ribosomal coordinates. This yielded a final curated dataset comprising 22,338 mHsp60:mHsp10 footballs, 6,130 mHsp60:mHsp10 half-footballs, 18,144 mHsp60 single-rings, 1,421 55S, and 3,647 39S mitochondrial ribosomes. Final sub-volumes were reconstructed in Warp at 4.6248 Å/pixel and refined in RELION 4.0. Particle poses were further optimized in M using successive geometric and CTF refinement steps, resulting in consensus maps at the following resolutions: 7.76 Å (mHsp60:mHsp10 football), 14.30 Å (mHsp60:mHsp10 half-football), 13.67 Å (mHsp60 single-ring), 22.89 Å (55S), and 15.92 Å (39S).

### Distance-based enrichment analysis

To assess the spatial enrichment of particles in the proximity of aggregates, a proximal region was defined as the volume within 50 nm of the nearest aggregate surface in 3D segmentations. For comparison, a distal region was defined to comprise an equivalent volume beginning from 50 nm distance to the aggregate and extending outwards. . The mitochondrial volume segmentation was used to compute the available volume within each region, and the Euclidean distance from each particle coordinate to the nearest aggregate voxel was calculated. The number of particles within the proximal and distal regions was normalized to their respective volumes to calculate the mean particle density per region and per tomogram. This allowed us to compare particle localization independently of the differing available space, enabling a direct assessment of particle enrichment near aggregates. These analyses were implemented in MuRePP (Multidimensional Representation of Particle Properties), a collection of MATLAB scripts for quantitative analysis of subtomogram averaging particle lists, compatible with RELION-style .star files (https://gitlab.gwdg.de/fernandez-busnadiego-lab/mitochondrial-proteostasis-cryo-et-analysis).

### Substrate protein-focused classification and analysis of mHsp60 single-rings

To investigate the occupancy states of the central cavity in mHsp60 single-rings, we applied focused classification using a cylindrical mask centered on the cavity while minimizing inclusion of surrounding mHsp60 density. Unbinned sub-volumes (2.3124 Å/pixel) were reconstructed in WARP following M-refinement and subjected to 3D classification in RELION 4.0 without alignment. Classification was performed with a regularization parameter of T=4, restricted to 25 Å resolution, and C7 symmetry imposed. Four classes were computed over 35 iterations using the focused mask. The resulting classes displayed distinct densities within the central cavity, localized either at the equatorial domain, the apical domain, or both. To validate the reproducibility of these classes, the classification procedure was independently repeated with identical parameters. Particles consistently assigned to the same class across both runs were grouped for downstream analysis. Four reproducible classes were identified: one featuring additional density in the equatorial region (3,015 particles), two with density in both the equatorial and apical regions (1,795 and 679 particles, respectively), and one with no additional cavity density (10,275 particles).

To assess potential conformational differences associated with SP binding, all particles from the three classes displaying additional densities were pooled and compared to the unoccupied class. A spherical mask was used to perform 3D refinement in RELION 4.0 followed by M-refinement. Atomic models of the ADP-bound, ATP-bound, and apo forms of the mHsp60 single-ring were docked into the resulting maps using rigid-body fitting in ChimeraX (Meng *et al*, 2023). The ADP-bound model consistently provided the best fit and was thus selected for further analysis of SP interaction.

Application of C7 symmetry enhanced the rotational signal and enabled localization of recurring SP interaction regions. Rigid-body docking of the ADP-bound model into each map allowed identification of contact sites between the additional density and the mHsp60 ring at contour levels corresponding to initial interaction. To control for potential artifacts introduced by symmetry, the classification was repeated in C1. Although the resulting classes exhibited high heterogeneity and did not fully converge, inspection of individual classes confirmed the interaction sites previously observed under C7 symmetry.

### Substrate protein-focused classification and analysis of mHsp60:mHsp10 half- and full-footballs

To investigate chamber occupancy in mHsp60:mHsp10 complexes, we employed a focused classification strategy adapted from (Wagner *et al*., 2024). Full mHsp60:mHsp10 football complexes were computationally split at the inter-ring interface, generating two mHsp60:mHsp10 half-footballs per complex. These artificially generated half-footballs were combined with *bona fide* half-footballs and jointly aligned using 3D refinement in RELION 4.0 followed by M-refinement. Unbinned sub-volumes were reconstructed in WARP at a pixel size of 2.3124 Å, and a cylindrical mask was designed to isolate the chamber interior while minimizing inclusion of surrounding mHsp60 density. To exclude unoccupied chambers, an initial 3D classification was performed in RELION 4.0 without alignment, using the chamber- focused mask, a regularization parameter of T=4, C7 symmetry, and a resolution limit of 30 Å over 35 iterations. The classification was repeated to assess reproducibility, and particles consistently assigned to the unoccupied class were designated as such. The remaining particles were pooled and subjected to a second round of classification under identical conditions but restricted to three classes only and a resolution limit of 25 Å to improve convergence. This round was also repeated independently, and particles stably classified in both runs were assigned to their respective class identities. This procedure yielded four major SP occupancy classes: one showing density localized to the equatorial region (20,279 particles), one with density primarily in the apical region (9,861 particles), one with density spanning both equatorial and apical regions (3,990 particles), and one unoccupied class (12,560 particles).

SP contact sites were identified following the same approach as used for mHsp60 single-rings. Briefly, the atomic model of the mHsp60:mHsp10 half-football was rigid body fitted into the STA density map upon application of C7 symmetry, and contour levels were adjusted to visualize the initial contact points between the additional density and the mHsp60:mHsp10 complex. To exclude potential artifacts introduced by imposing symmetry, the entire classification was repeated under C1 symmetry.

To assess whether specific combinations of SP localization within a full mHsp60:mHsp10 football complex occurred more frequently than expected by chance, we modeled a null distribution of expected pairings. The relative abundance of each class within the full football set was treated as its marginal probability, and the expected frequency of all possible class pairings was calculated under the assumption of random association (i.e., as the product of their marginal probabilities). The 18,845 mHsp60:mHsp10 football complexes detected in the complete dataset was used as total.

### Visualizations

All 3D renderings of segmentations, cryo-EM maps, and composite views were generated using ChimeraX with the ArtiaX plugin (Ermel *et al*, 2022).

### Statistics

Statistical analyses for pairwise comparisons of membrane densities (Figure 1C, D) and particle densities (Figure 2B, E, H) were performed using the nonparametric Wilcoxon rank-sum test. For the cell viability assay (Figure S 1B), significance of pairwise comparisons was assessed using a two-sample t-test. Significance was indicated as follows: p<0.05 as *, p<0.01 as ** and p<0.001 as ***.

