## Supplementary information for "Structural remodeling of the mitochondrial protein biogenesis machinery under proteostatic stress"

Figure S1

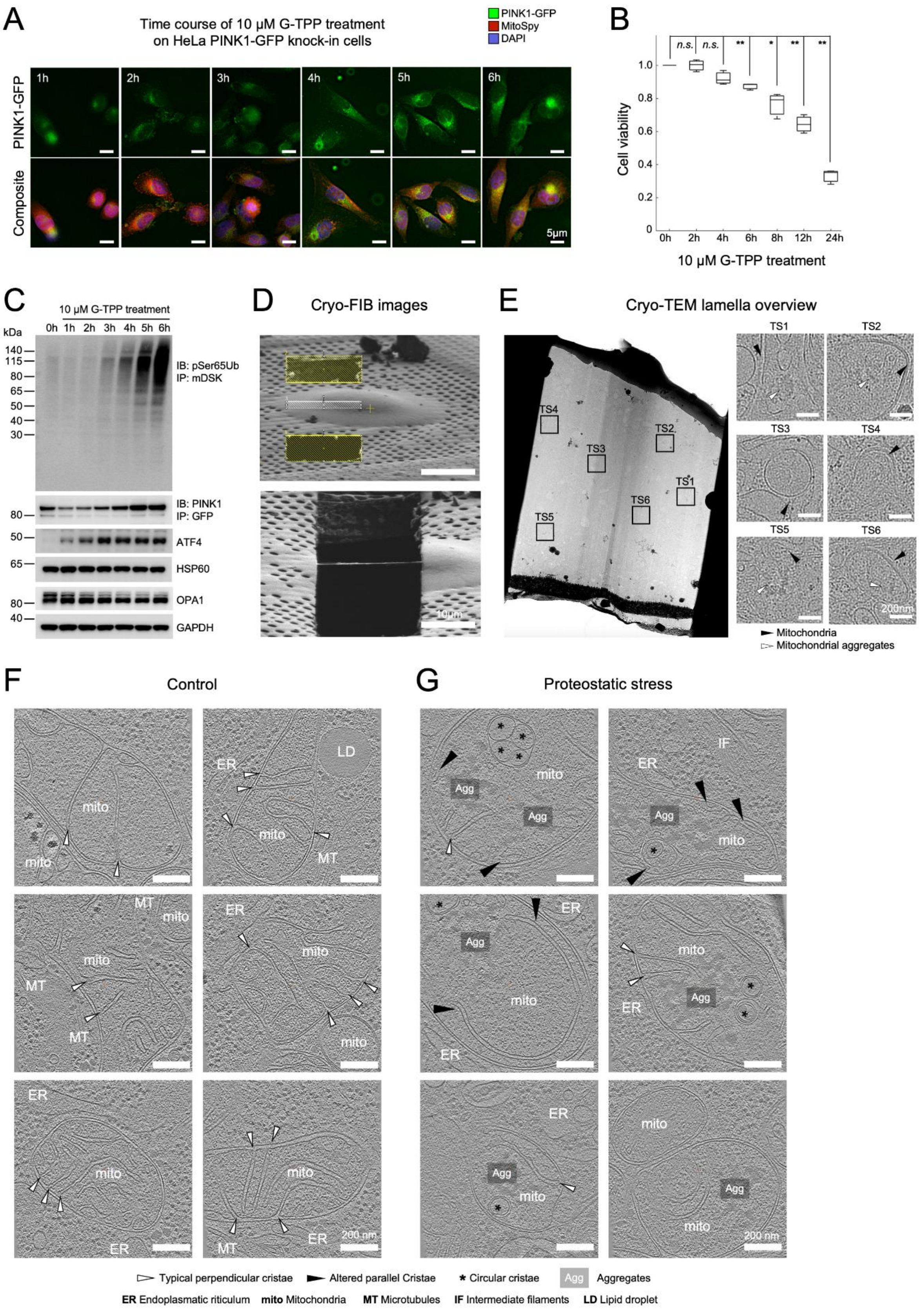
A
Time course of 10 μM G-TPP treatment on HeLa PINK1-GFP knock-in cells
PINK1-GFP
MitoSpy
DAPI
1h
2h
3h
4h
5h
6h
PINK1-GFP
Composite
5μm
B
n.s.
n.s.
**
*
**
**
Cell viability
0h
2h
4h
6h
8h
12h
24h
10 μM G-TPP treatment
C
10 μM G-TPP treatment
kDa
IB: pSer65Ub
IP: mDSK
IB: PINK1
IP: GFP
ATF4
HSP60
OPA1
GAPDH
D
Cryo-FIB images
10μm
E
Cryo-TEM lamella overview
TS1
TS2
TS3
TS4
TS5
TS6
200nm
Mitochondria
Mitochondrial aggregates
F
Control
G
Proteostatic stress
ER
LD
mito
MT
IF
Agg
200 nm
Typical perpendicular cristae
Altered parallel Cristae
Circular cristae
Aggregates
ER Endoplasmatic riticulum
mito Mitochondria
MT Microtubules
IF Intermediate filaments
LD Lipid droplet

**Figure S 1: G-TPP treatment induces proteostatic stress in HeLa cells.** (**A**) Fluorescence microscopy of PINK1-GFP HeLa cells at time intervals following 10 µM G-TPP treatment. Increasing co-localization of PINK1-GFP with the mitochondrial marker MitoSpy indicates PINK1 stabilization at mitochondria upon proteostatic stress. (**B**) Quantification of cell viability as a function of time following 10 µM G-TPP treatment. Statistical significance of pairwise comparisons was assessed using a two-sample t-test and indicated by: n.s. ($p > 0.05$), * ($p < 0.05$) and ** ($p < 0.01$). N = 3 independent experiments. (**C**) Western blot analysis of the effects of G-TPP treatment on PINK1 accumulation and activation, measured by increased levels of ubiquitin phosphorylation at serine 65 (pSer65Ub). OPA1 band shifts and ATF4 activation serve as additional indicators of mitochondrial stress. (**D**) Ion beam-induced secondary electron images of a HeLa cell vitrified on an EM grid, shown during (top) and after (bottom) cryo-FIB milling. (**E**) Left: cryo-TEM image of a cryo-FIB-milled lamella from a vitrified HeLa cell that underwent 10 µM G-TPP treatment. Boxes mark regions selected for tomographic tilt series (TS) acquisition. Right: Magnified views of the regions of tomographic data acquisition highlighting mitochondria (black arrowheads). In some cases, mitochondrial aggregates (white arrowheads) were readily distinguishable. (**F–G**) Gallery of tomographic slices from untreated control (F) and G-TPP–treated cells under proteostatic stress (G). Cellular structures are annotated in the slices according to the legend at the bottom of the figure.

Figure S2

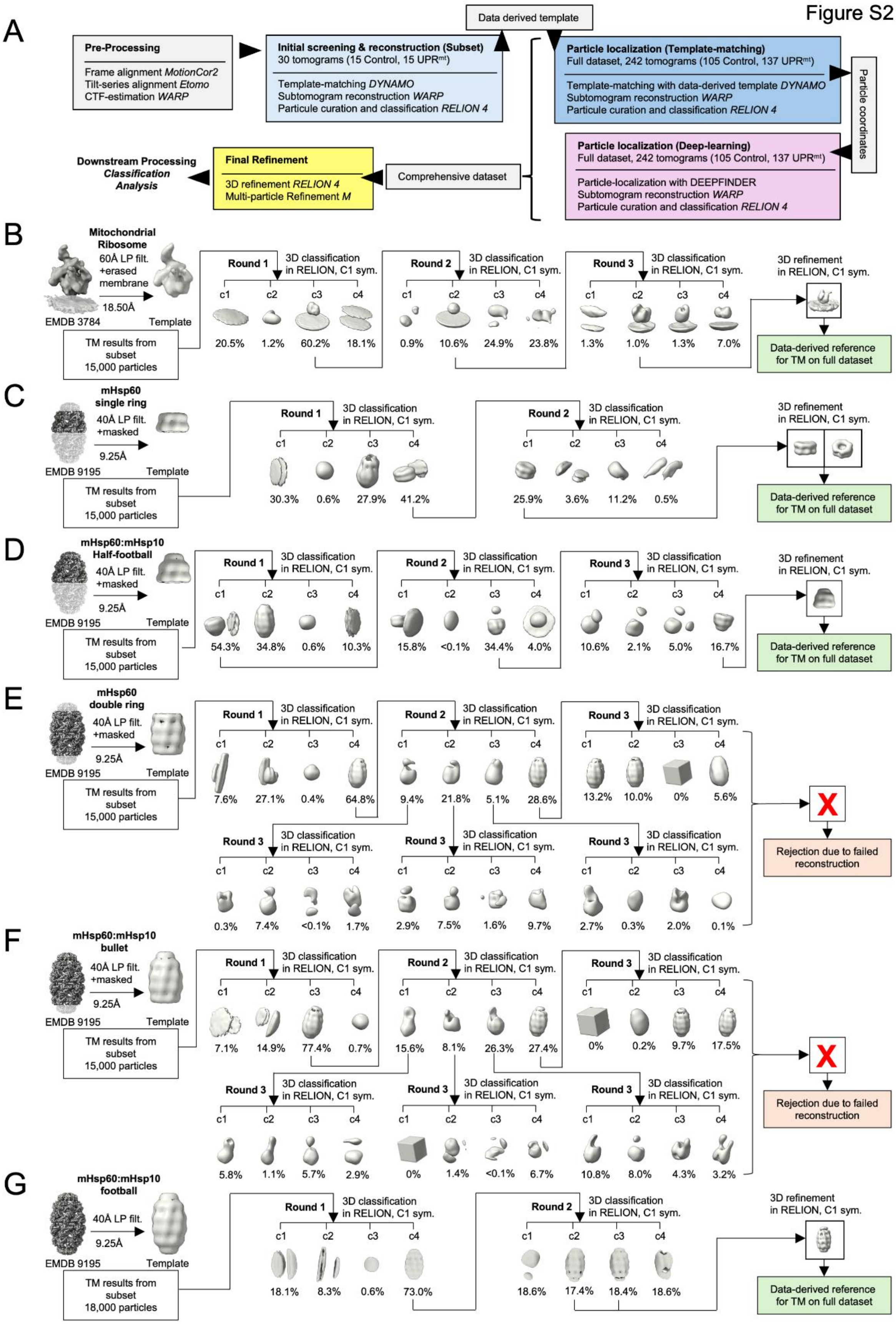
A
Pre-Processing
Frame alignment MotionCor2
Tilt-series alignment Etomo
CTF-estimation WARP
Initial screening & reconstruction (Subset)
30 tomograms (15 Control, 15 UPRmt)
Template-matching DYNAMO
Subtomogram reconstruction WARP
Particule curation and classification RELION 4
Data derived template
Particle localization (Template-matching)
Full dataset, 242 tomograms (105 Control, 137 UPRmt)
Template-matching with data-derived template DYNAMO
Subtomogram reconstruction WARP
Particule curation and classification RELION 4
Particle coordinates
Particle localization (Deep-learning)
Full dataset, 242 tomograms (105 Control, 137 UPRmt)
Particle-localization with DEEPFINDER
Subtomogram reconstruction WARP
Particule curation and classification RELION 4
Comprehensive dataset
Final Refinement
3D refinement RELION 4
Multi-particle Refinement M
Downstream Processing
Classification
Analysis
B
Mitochondrial Ribosome
60Å LP filt. +erased membrane
18.50Å
EMDB 3784
Template
TM results from subset 15,000 particles
Round 1
3D classification in RELION, C1 sym.
c1 c2 c3 c4
20.5% 1.2% 60.2% 18.1%
Round 2
3D classification in RELION, C1 sym.
c1 c2 c3 c4
0.9% 10.6% 24.9% 23.8%
Round 3
3D classification in RELION, C1 sym.
c1 c2 c3 c4
1.3% 1.0% 1.3% 7.0%
3D refinement in RELION, C1 sym.
Data-derived reference for TM on full dataset
C
mHsp60 single ring
40Å LP filt. +masked
9.25Å
EMDB 9195
Template
TM results from subset 15,000 particles
Round 1
3D classification in RELION, C1 sym.
c1 c2 c3 c4
30.3% 0.6% 27.9% 41.2%
Round 2
3D classification in RELION, C1 sym.
c1 c2 c3 c4
25.9% 3.6% 11.2% 0.5%
3D refinement in RELION, C1 sym.
Data-derived reference for TM on full dataset
D
mHsp60:mHsp10 Half-football
40Å LP filt. +masked
9.25Å
EMDB 9195
Template
TM results from subset 15,000 particles
Round 1
3D classification in RELION, C1 sym.
c1 c2 c3 c4
54.3% 34.8% 0.6% 10.3%
Round 2
3D classification in RELION, C1 sym.
c1 c2 c3 c4
15.8% <0.1% 34.4% 4.0%
Round 3
3D classification in RELION, C1 sym.
c1 c2 c3 c4
10.6% 2.1% 5.0% 16.7%
3D refinement in RELION, C1 sym.
Data-derived reference for TM on full dataset
E
mHsp60 double ring
40Å LP filt. +masked
9.25Å
EMDB 9195
Template
TM results from subset 15,000 particles
Round 1
3D classification in RELION, C1 sym.
c1 c2 c3 c4
7.6% 27.1% 0.4% 64.8%
Round 2
3D classification in RELION, C1 sym.
c1 c2 c3 c4
9.4% 21.8% 5.1% 28.6%
Round 3
3D classification in RELION, C1 sym.
c1 c2 c3 c4
13.2% 10.0% 0% 5.6%
Round 3
3D classification in RELION, C1 sym.
c1 c2 c3 c4
0.3% 7.4% <0.1% 1.7%
Round 3
3D classification in RELION, C1 sym.
c1 c2 c3 c4
2.9% 7.5% 1.6% 9.7%
Round 3
3D classification in RELION, C1 sym.
c1 c2 c3 c4
2.7% 0.3% 2.0% 0.1%
X
Rejection due to failed reconstruction
F
mHsp60:mHsp10 bullet
40Å LP filt. +masked
9.25Å
EMDB 9195
Template
TM results from subset 15,000 particles
Round 1
3D classification in RELION, C1 sym.
c1 c2 c3 c4
7.1% 14.9% 77.4% 0.7%
Round 2
3D classification in RELION, C1 sym.
c1 c2 c3 c4
15.6% 8.1% 26.3% 27.4%
Round 3
3D classification in RELION, C1 sym.
c1 c2 c3 c4
0% 0.2% 9.7% 17.5%
Round 3
3D classification in RELION, C1 sym.
c1 c2 c3 c4
5.8% 1.1% 5.7% 2.9%
Round 3
3D classification in RELION, C1 sym.
c1 c2 c3 c4
0% 1.4% <0.1% 6.7%
Round 3
3D classification in RELION, C1 sym.
c1 c2 c3 c4
10.8% 8.0% 4.3% 3.2%
X
Rejection due to failed reconstruction
G
mHsp60:mHsp10 football
40Å LP filt.
9.25Å
EMDB 9195
Template
TM results from subset 18,000 particles
Round 1
3D classification in RELION, C1 sym.
c1 c2 c3 c4
18.1% 8.3% 0.6% 73.0%
Round 2
3D classification in RELION, C1 sym.
c1 c2 c3 c4
18.6% 17.4% 18.4% 18.6%
3D refinement in RELION, C1 sym.
Data-derived reference for TM on full dataset

**Figure S 2: Cryo-ET particle picking workflow and localization of mitochondrial protein complexes by template matching.** (**A**) Schematic overview of the processing strategy. After initial preprocessing, mitochondrial complexes were identified and their relative abundance assessed by template matching in DYNAMO and subtomogram averaging in RELION 4.0 on a subset of data. Following successful reconstruction, the resulting density maps were used as templates for template matching on the full datasets. Curated particle coordinates were then input into DEEPFINDER for deep-learning–based particle detection to improve accuracy and coverage. The combined datasets from both approaches were refined in RELION 4.0 and M to generate final density maps and particle orientations for downstream analyses. (**B–G**) Classification steps for the mitochondrial ribosome and mHsp60 complexes during initial screening. To detect the presence of various mHsp60 complexes, the single-particle cryo-EM density map of a human mHsp60:mHsp10 football complex (EMDB 9195) was masked to generate references of the different complexes for initial classification. Upon masking, the reference maps were then low-pass filtered to 60 Å (B) or 40 Å (C-G) resolution and rescaled to 18.5 Å (B) or 9.25 Å (C-G) pixel size, to match the dimensions of the tomogram in the respective binning. All classification steps were performed without imposing symmetry. Abbreviations: C1 sym, C1 symmetry; LP filt, low-pass filtered; TM, template matching.

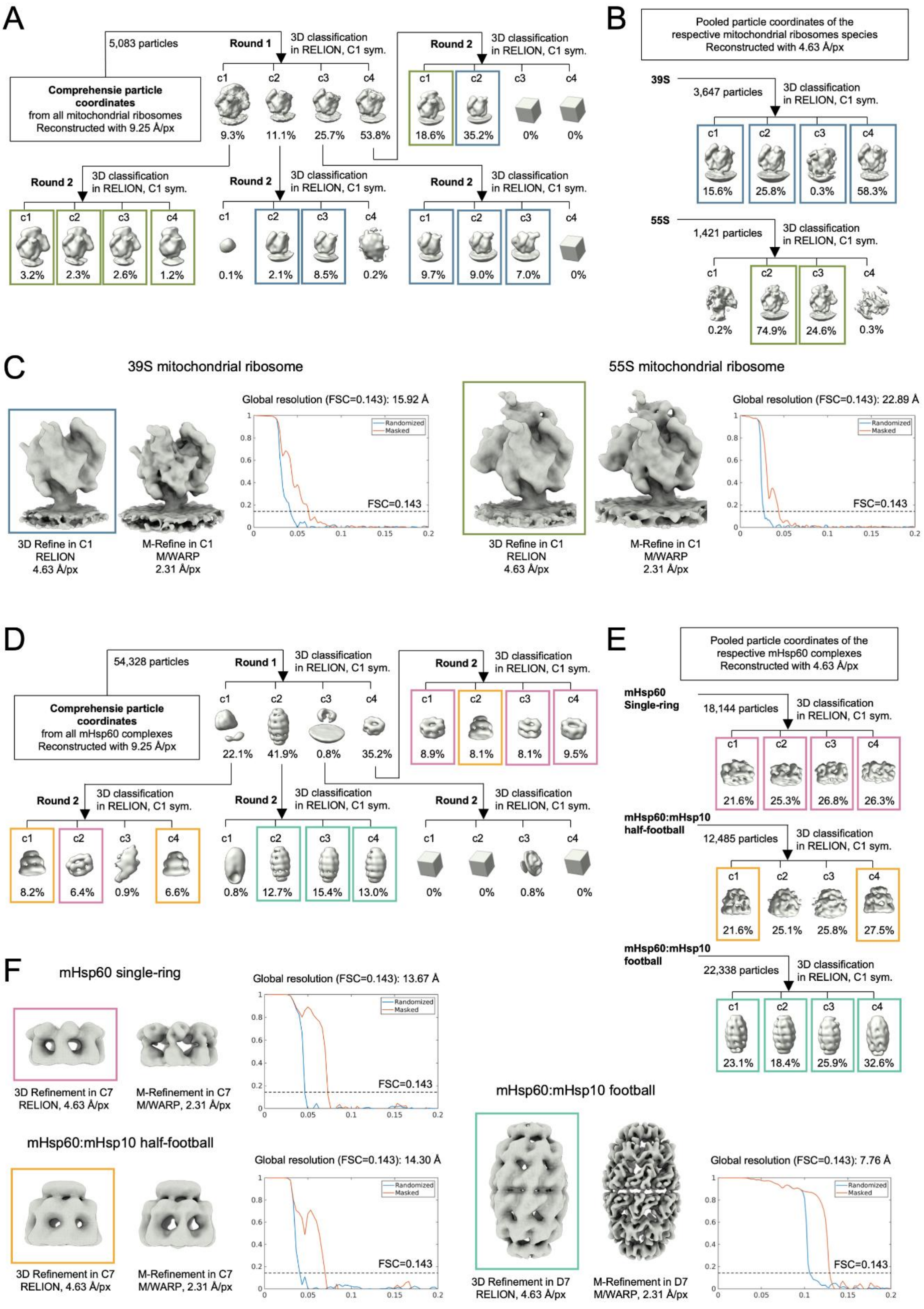
A
5,083 particles
Round 1
3D classification in RELION, C1 sym.
Comprehensie particle coordinates from all mitochondrial ribosomes Reconstructed with 9.25 Å/px
c1 c2 c3 c4
9.3% 11.1% 25.7% 53.8%
Round 2
3D classification in RELION, C1 sym.
18.6% 35.2% 0% 0%
3.2% 2.3% 2.6% 1.2%
0.1% 2.1% 8.5% 0.2%
9.7% 9.0% 7.0% 0%
B
Pooled particle coordinates of the respective mitochondrial ribosomes species Reconstructed with 4.63 Å/px
39S
3,647 particles
15.6% 25.8% 0.3% 58.3%
55S
1,421 particles
0.2% 74.9% 24.6% 0.3%
C
39S mitochondrial ribosome
Global resolution (FSC=0.143): 15.92 Å
3D Refine in C1 RELION 4.63 Å/px
M-Refine in C1 M/WARP 2.31 Å/px
55S mitochondrial ribosome
Global resolution (FSC=0.143): 22.89 Å
Randomized
Masked
FSC=0.143
D
54,328 particles
Comprehensie particle coordinates from all mHsp60 complexes Reconstructed with 9.25 Å/px
22.1% 41.9% 0.8% 35.2%
8.9% 8.1% 8.1% 9.5%
8.2% 6.4% 0.9% 6.6%
0.8% 12.7% 15.4% 13.0%
0% 0% 0.8% 0%
E
Pooled particle coordinates of the respective mHsp60 complexes Reconstructed with 4.63 Å/px
mHsp60 Single-ring
18,144 particles
21.6% 25.3% 26.8% 26.3%
mHsp60:mHsp10 half-football
12,485 particles
21.6% 25.1% 25.8% 27.5%
mHsp60:mHsp10 football
22,338 particles
23.1% 18.4% 25.9% 32.6%
F
mHsp60 single-ring
Global resolution (FSC=0.143): 13.67 Å
3D Refinement in C7 RELION, 4.63 Å/px
M-Refinement in C7 M/WARP, 2.31 Å/px
mHsp60:mHsp10 half-football
Global resolution (FSC=0.143): 14.30 Å
mHsp60:mHsp10 football
Global resolution (FSC=0.143): 7.76 Å
3D Refinement in D7 RELION, 4.63 Å/px
M-Refinement in D7 M/WARP, 2.31 Å/px

**Figure S 3: *In situ* structural determination of mitochondrial ribosomes and mHsp60 complexes.** (**A, D**) Curated and distance-filtered particle coordinates for mitochondrial ribosomes (A) and mHsp60 complexes (D). Particles were pooled and reclassified at bin 4 to identify the different assemblies. (**B, E**) Each assembly was further classified at bin 2 to remove remaining false positives and obtain the final particle sets used for refinement. (**C, F**) Final subtomogram averaging density maps were obtained after refinement at bin 2 in RELION 4.0, followed by multi-particle refinement at bin 1 using M. Abbreviations: C1 sym, C1 symmetry; FSC, Fourier shell correlation; px, pixel.

Figure S4

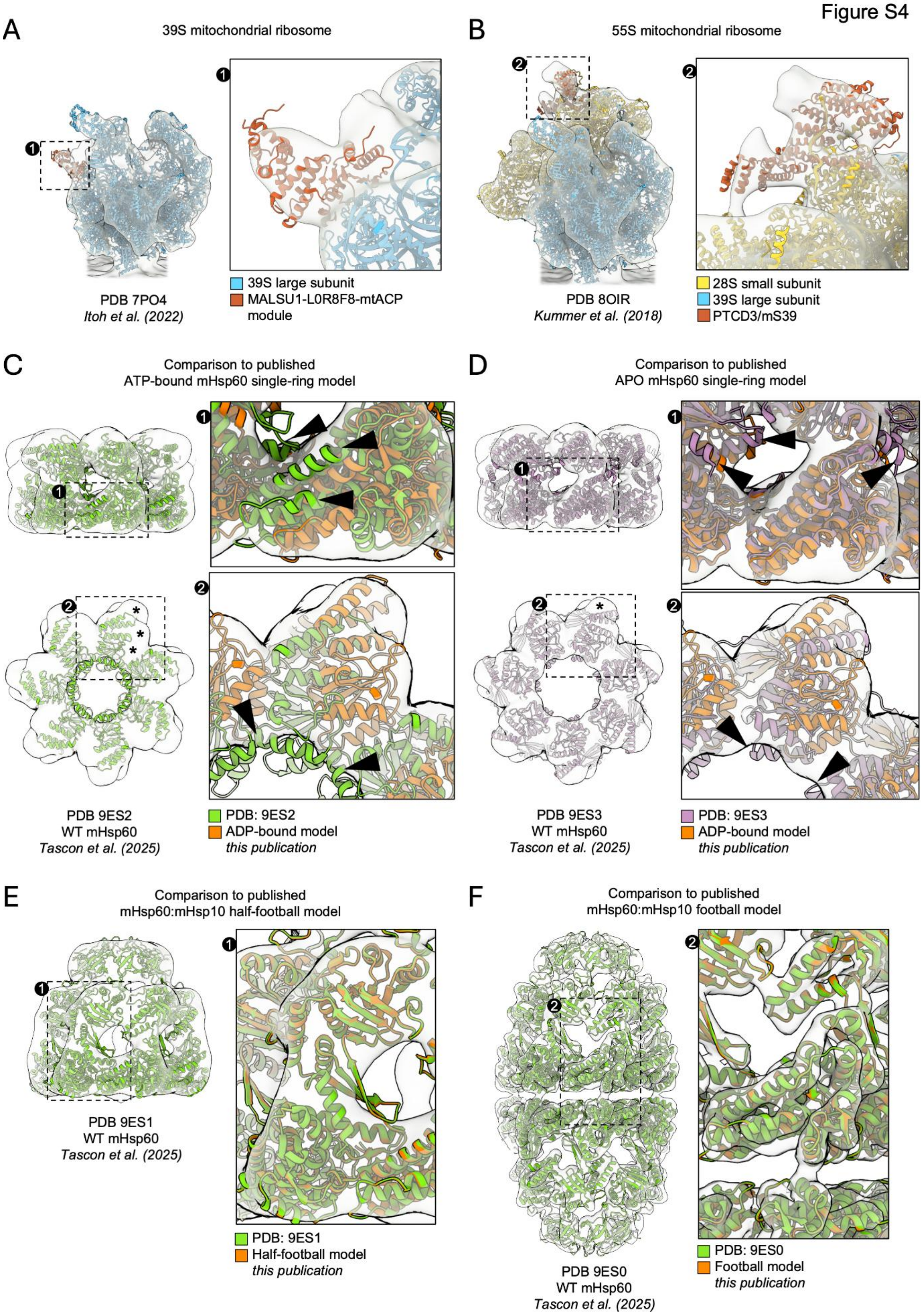
A
39S mitochondrial ribosome
1
PDB 7PO4
Itoh et al. (2022)
39S large subunit
MALSU1-L0R8F8-mtACP module
B
55S mitochondrial ribosome
2
PDB 8OIR
Kummer et al. (2018)
28S small subunit
39S large subunit
PTCD3/mS39
C
Comparison to published
ATP-bound mHsp60 single-ring model
PDB 9ES2
WT mHsp60
Tascon et al. (2025)
PDB: 9ES2
ADP-bound model
this publication
D
Comparison to published
APO mHsp60 single-ring model
PDB 9ES3
WT mHsp60
Tascon et al. (2025)
PDB: 9ES3
ADP-bound model
this publication
E
Comparison to published
mHsp60:mHsp10 half-football model
PDB 9ES1
WT mHsp60
Tascon et al. (2025)
PDB: 9ES1
Half-football model
this publication
F
Comparison to published
mHsp60:mHsp10 football model
PDB 9ES0
WT mHsp60
Tascon et al. (2025)
PDB: 9ES0
Football model
this publication

**Figure S 4: Docking of atomic models into subtomogram averaging density maps.** In all cases, *in situ* subtomogram averaging map are shown semitransparent. (**A**) Docking of the atomic model of a 39S mitochondrial ribosome assembly intermediate (PDB 7PO4). The inset highlights density consistent with the binding of the MALSU1–L0R8F8–mtACP module (Hillen *et al*, 2021; Lavdovskaia *et al*, 2024; Rebelo-Guiomar *et al*, 2022). (**B**) Docking of the atomic model of a fully assembled 55S mitochondrial ribosome (PDB 8OIR). The inset indicates density consistent with the binding of the RNA-binding and translation-regulating PTCD3/mS39 subunit (Kummer *et al*, 2018). (**C–F**) Docking of atomic models of WT mHsp60 complexes derived from single-particle cryo-EM into subtomogram averaging maps determined *in situ* using cryo-ET. We compare the cryo-EM structures determined in this study (orange) with published structures of mHSp60 complexes (green, purple). For clarity, comparisons are only shown with structures from (Tascon *et al*, 2025), which are representative of most other published mHps60 structures (Table S 1). (C, D) Comparison of the fit into subtomogram maps of ATP-bound (C) and apo (D) mHsp60 single-ring structures against ADP-bound mHsp60. Regions where the docked ATP and apo structures do not fit well the subtomogram density map are indicated by black arrowheads (structures extending beyond the map) and stars (densities in the map unaccounted for by the structures). (E, F) Overlays of published mHsp60:mHsp10 structures indicate only minimal variations compared to the ones determined in this study.

Figure S5

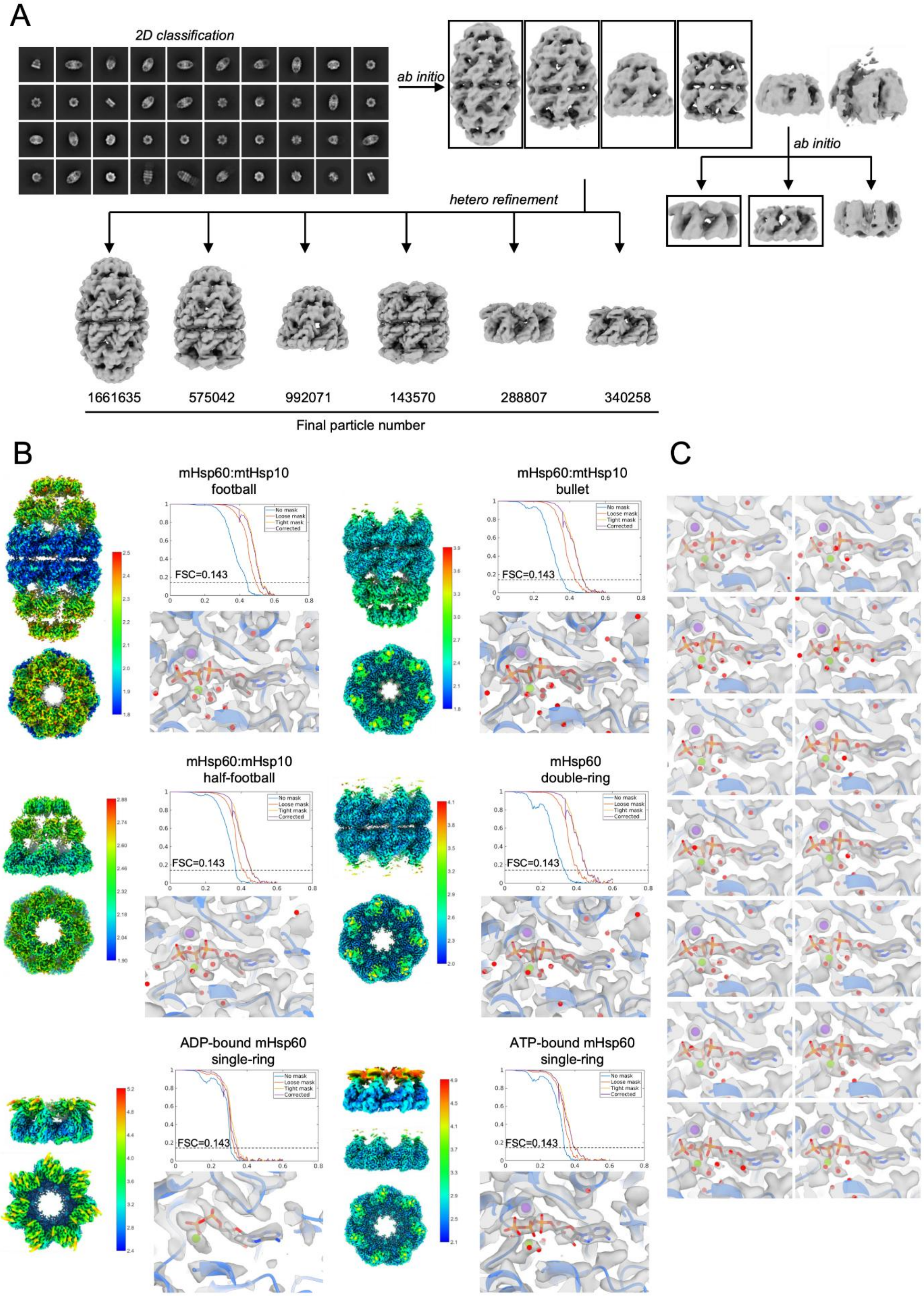
A
2D classification
ab initio
ab initio
hetero refinement
1661635
575042
992071
143570
288807
340258
Final particle number
B
mHsp60:mtHsp10
football
mHsp60:mtHsp10
bullet
mHsp60:mHsp10
half-football
mHsp60
double-ring
ADP-bound mHsp60
single-ring
ATP-bound mHsp60
single-ring
FSC=0.143
No mask
Loose mask
Tight mask
Corrected
C

**Figure S 5: Structures of mHsp60 complexes determined *in vitro* by single-particle cryo-EM.** (**A**) Cryo-EM data processing workflow for mHsp60 complexes. All datasets were processed independently, starting with particle picking, iterative 2D classification, and *ab initio* reconstruction. An additional *ab initio* reconstruction was performed specifically for particles contributing to single-ring classes. Boxes mark the *ab initio* volumes used for heterogeneous refinement. (**B**) Cryo-EM maps of ATP-bound mHsp60:mHsp10 football (1.92 Å nominal resolution), ATP-bound mHsp60:mHsp10 half-football (2.19 Å nominal resolution), ATP-bound mHsp60:mHsp10 bullet (2.1 Å nominal resolution), ADP-bound mHsp60 single-ring (2.91 Å nominal resolution), ATP-bound mHsp60 double ring (2.18 Å nominal resolution), and ATP-bound mHsp60 single-ring (2.50 Å nominal resolution) colored as a function of local resolution displayed as side and top views. FSC curves and detailed views of the nucleotide-binding sites for each species are also shown, displaying overlays of the density map (semitransparent) fitted with atomic models. The protein is depicted as blue ribbon, nucleotides as sticks, and $Mg^{2+}$ ions (green), $K^{+}$ ions (purple) and water molecules (red) as spheres. Cryo-EM maps were refined with D7 symmetry for the football and double-ring complexes, and C7 symmetry for the half-football, bullet, single-ring assemblies. For the ATP-bound mHsp60 single-ring, the map is also shown upon gaussian-filtering (top) to enable visualization of the highly-dynamic apical domains. (**C**) Nucleotide-binding sites of all 14 subunits of the ATP-bound mHsp60:mHsp10 football complex obtained by refinement without imposing symmetry (C1 symmetry). Abbreviations: FSC, Fourier shell correlation.

Figure S6

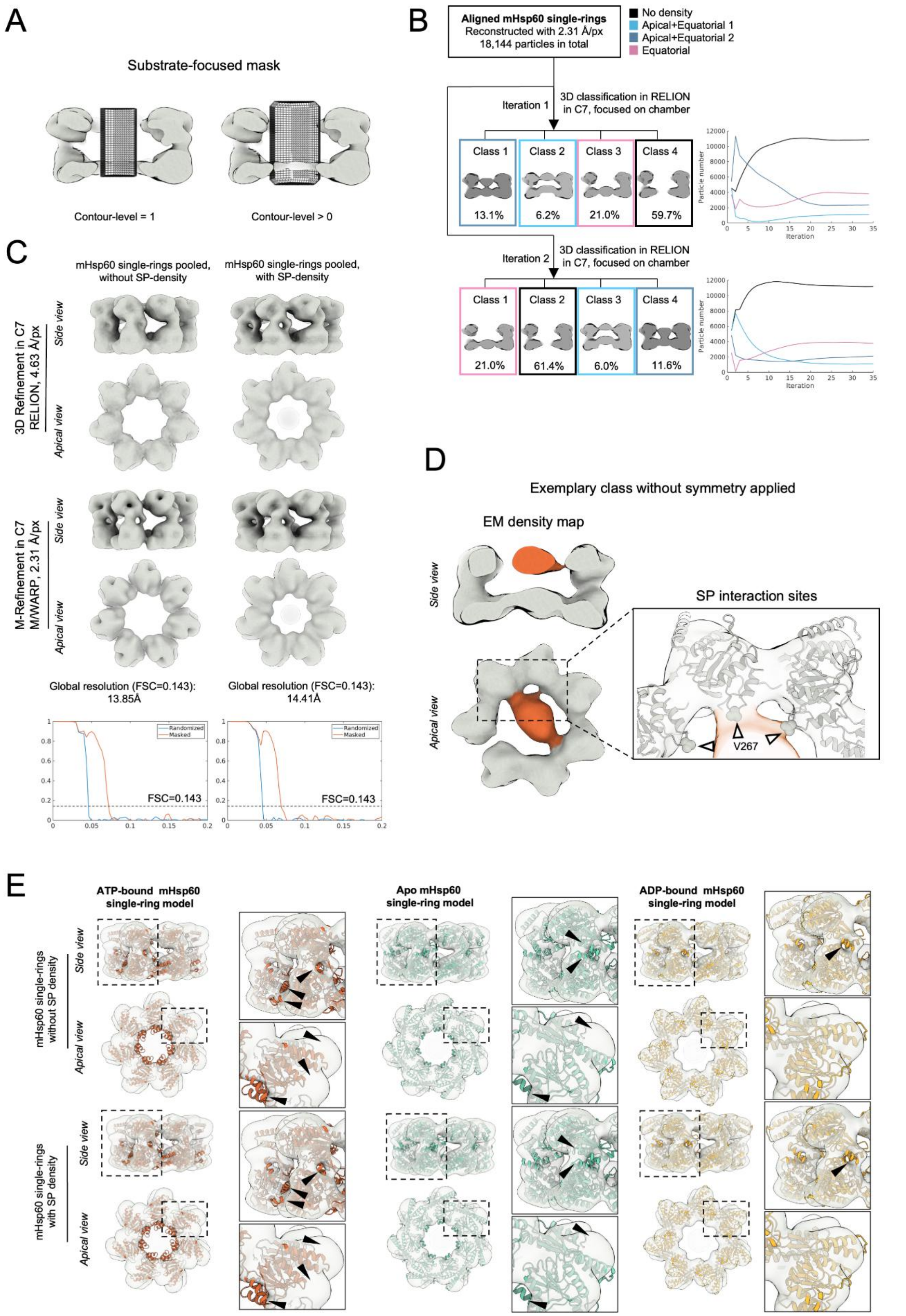
A
Substrate-focused mask
Contour-level = 1
Contour-level > 0
B
Aligned mHsp60 single-rings
Reconstructed with 2.31 Å/px
18,144 particles in total
No density
Apical+Equatorial 1
Apical+Equatorial 2
Equatorial
Iteration 1
3D classification in RELION in C7, focused on chamber
Class 1
Class 2
Class 3
Class 4
13.1%
6.2%
21.0%
59.7%
Particle number
Iteration
Iteration 2
3D classification in RELION in C7, focused on chamber
21.0%
61.4%
6.0%
11.6%
C
mHsp60 single-rings pooled, without SP-density
mHsp60 single-rings pooled, with SP-density
3D Refinement in C7 RELION, 4.63 Å/px
Side view
Apical view
M-Refinement in C7 M/WARP, 2.31 Å/px
Global resolution (FSC=0.143): 13.85Å
Global resolution (FSC=0.143): 14.41Å
Randomized
Masked
FSC=0.143
D
Exemplary class without symmetry applied
EM density map
SP interaction sites
V267
E
ATP-bound mHsp60 single-ring model
Apo mHsp60 single-ring model
ADP-bound mHsp60 single-ring model
mHsp60 single-rings without SP density
mHsp60 single-rings with SP density

**Figure S 6: Substrate protein classification in mHsp60 single-ring complexes.** (**A**) Mask of the central cavity used for focused 3D classification shown at different contour levels. The rendering at contour level = 1 represents the mask without the soft edge, while levels >0 visualize the full extent of the mask including the soft-edged boundary. (**B**) Classification results (left) and class occupancies over classification iterations (right) for two independent runs. (**C**) Independent refinements of the "No density" class versus SP-bound classes to assess potential structural differences. (**D**) Exemplary SP-bound class obtained from focused chamber-focused 3D classification in C1. The inset highlights the interactions between the SP density and V267 residues of mHsp60 (shown as spheres and marked by white arrowheads), as also observed in the classification using C7 symmetry (Figure 5B). (**E**) Docking of mHsp60 single-ring atomic models in different nucleotide states into the final subtomogram averaging maps from (C). Insets show magnified regions of interest, with black arrowheads indicating discrepancies between the model and the density map. Abbreviations: FSC, Fourier shell correlation; px, pixel.

Figure S7

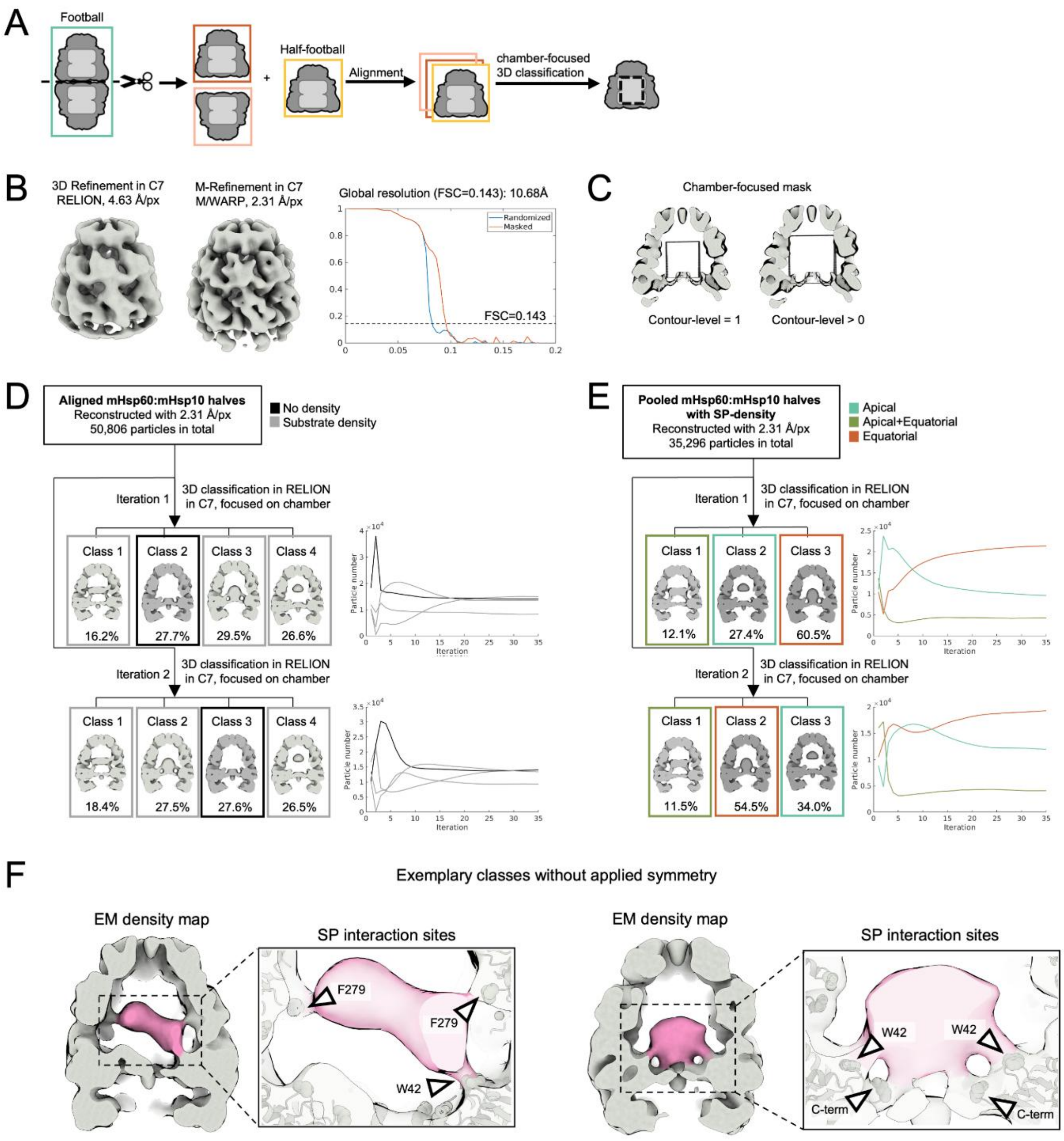

**Figure S 7: Substrate protein classification in mHsp60:mHsp10 complexes.** (**A**) Schematic overview of the processing workflow. mHsp60:mHsp10 football complexes were computationally bisected and their halves were aligned to *bona fide* mHsp60:mHsp10 half-football complexes prior to classification. (**B**) Refinement results for pooled mHsp60:mHsp10 half-maps including both *bona fide* mHsp60:mHsp10 half-footballs and bisected footballs. (**C**) Mask of the central cavity at different contour levels used for focused 3D classification. The rendering at contour level = 1 represents the mask without the soft edge, while levels >0 visualize the full extent of the mask including the soft-edged boundary. (**D**) First round of classification identifying empty chambers. Class averages (left) and class occupancies across iterations (right) are shown for two independent runs. (**E**) Second round of classification identifying different SP localization patterns. Class averages (left) and class occupancies across iterations (right) shown for two independent runs. (**F**) Examples of two different SP-bound classes obtained via focused 3D classification in C1. Insets highlight mHsp60 residues interacting with SPs are shown as spheres and indicated by arrowheads. These interaction sites were also identified using classification with applied symmetry (Figure 6B).

**Table S 1: RMSD comparison between published single-particle cryo-EM structures of mHsp60 complexes and those determined in this study**. Calculations were performed using the Matchmaker tool within ChimeraX (Goddard *et al*, 2018). The structures of WT mHsp60 complexes recently published by (Tascon *et al.*, 2025) were used as reference (9ES0, 9ES1, 9ES2, 9ES3). Only one ring of the ATP-bound mHsp60 double ring 9ES2 model was used for the calculations. The following structures of the V72I mHsp60 mutant were published by (Braxton *et al*, 2024): 8G7N (ATP-bound football), 8G7O (ATP-bound half-football), 8G7M (ATP-bound single-ring) and 8G7K (apo single-ring). The following structures of WT mHsp60 were published by (Gomez-Llorente *et al*, 2020): 6MRC (ADP-bound football), 6MRD (ADP-bound half-football). The 7AZP of WT mHsp60 apo single ring was published by (Klebl *et al*, 2021). The 7L7S of WT mHsp60 apo single ring was published by (Wang & Chen, 2021).

| Reference model | Matching model | RMSD (all pairs) |
|---|---|---|
| 9ES0 (ATP football) | ATP football (this study) | 0.250 Å |
| (Tascon et al., 2025) | 8G7N | 0.729 Å |
| | 6MRC | 0.471 Å |
| 9ES1 (ATP half-football) | ATP half-football (this study) | 0.324 Å |
| (Tascon et al., 2025) | 8G7O | 0.780 Å |
| | 6MRD | 0.642 Å |
| 9ES2 (ATP double-ring) | ATP single-ring (this study) | 0.816 Å |
| (Tascon et al., 2025) | ADP single-ring (this study) | 2.658 Å |
| | 8G7M | 2.844 Å |
| 9ES3 (Apo single-ring) | ATP single-ring (this study) | 3.883 Å |
| (Tascon et al., 2025) | ADP single-ring (this study) | 7.289 Å |
| | 8G7K | 1.327 Å |
| | 7AZP | 0.545 Å |
| | 7L7S | 1.036 Å |

**Table S 2**: **Cryo-EM data collection, refinement and validation statistics**

| | ATP-bound mHsp60$_{14}$-mHsp1014$_{14}$ (EMDB 54898) (PDB 9SHG) | ATP-bound mHsp60$_{14}$-mHsp10$_{7}$ (EMDB 54899) (PDB 9SHH) | ATP-bound mHsp60$_{7}$-mHsp10$_{7}$ (EMDB 54990) (PDB 9SHI) |
|---|---|---|---|
| **Data collection and processing** | | | |
| Magnification | 105000 | 105000 | 105000 |
| Voltage (kV) | 300 | 300 | 300 |
| Electron exposure (e–/Å$^2$) | 50 | 50 | 50 |
| Defocus range (μm) | 0.8-1.6 | 0.8-1.6 | 0.8-1.6 |
| Pixel size (Å) | 0.8238 | 0.8238 | 0.8238 |
| Symmetry imposed | D7 | C7 | D7 |
| Initial particle images (no.) | 12338206 | 12338206 | 12338206 |
| Final particle images (no.) | 1661635 | 575042 | 992071 |
| Map resolution (Å) | 1.91 | 2.11 | 2.19 |
| FSC threshold | 0.143 | 0.143 | 0.143 |
| Map resolution range (Å) | 1.845 - 20.712 | 1.881 - 30.220 | 1.818 - 25.245 |
| **Refinement** | | | |
| Initial model used (PDB code) | 9ES0 | 9ES1/9ES2 | 9ES1 |
| Model resolution (Å) | 2.0 | 2.2 | 2.5 |
| FSC threshold | 0.5 | 0.5 | 0.5 |
| Map sharpening *B* factor (Å$^2$) | -60.4 | -60.3 | -75.5 |
| Model composition | | | |
| Non-hydrogen atoms | 68854 | 63040 | 34531 |
| Protein residues | 8778 | 8078 | 4389 |
| Ligands | 42 | 42 | 21 |
| *B* factors (Å$^2$) | | | |
| Protein | 43.14 | 69.33 | 47.69 |
| Ligand | 11.32 | 17.88 | 18.59 |
| R.m.s. deviations | | | |
| Bond lengths (Å) | 0.004 | 0.004 | 0.004 |
| Bond angles (°) | 0.680 | 0.676 | 0.636 |
| Validation | | | |
| MolProbity score | 1.05 | 1.20 | 1.10 |
| Clashscore | 2.65 | 4.21 | 3.13 |
| Poor rotamers (%) | 0.00 | 0.00 | 0.00 |
| Ramachandran plot | | | |
| Favored (%) | 98.73 | 98.36 | 98.62 |
| Allowed (%) | 1.27 | 1.64 | 1.38 |
| Disallowed (%) | 0.00 | 0.00 | 0.00 |

| | ATP-bound $mHsp60_{14}$ (EMDB 54991) (PDB 9SHJ) | ADP-bound $mHsp60_{7}$ (EMDB 54992) (PDB 9SHK) | ATP-bound $mHsp60_{7}$ (EMDB 54993) (PDB 9SHGL) |
|---|---|---|---|
| **Data collection and processing** | | | |
| Magnification | 105000 | 105000 | 105000 |
| Voltage (kV) | 300 | 300 | 300 |
| Electron exposure (e–/Å$^2$) | 50 | 50 | 50 |
| Defocus range (μm) | 0.8-1.6 | 0.8-1.6 | 0.8-1.6 |
| Pixel size (Å) | 0.8238 | 0.8238 | 0.8238 |
| Symmetry imposed | D7 | C7 | C7 |
| Initial particle images (no.) | 12338206 | 12338206 | 12338206 |
| Final particle images (no.) | 143570 | 288807 | 340258 |
| Map resolution (Å) | 2.18 | 2.91 | 2.50 |
| FSC threshold | 0.143 | 0.143 | 0.143 |
| Map resolution range (Å) | 1.978 - 34.455 | 1.768 - 9.351 | 2.147 - 24.030 |
| **Refinement** | | | |
| Initial model used (PDB code) | 9ES2 | 9ES1 | 9ES2 |
| Model resolution (Å) | 2.3 | 3.3 | 2.8 |
| FSC threshold | 0.5 | 0.5 | 0.5 |
| Map sharpening *B* factor (Å$^2$) | -58.0 | -106.0 | -97.6 |
| Model composition | | | |
| Non-hydrogen atoms | 56446 | 27629 | 28015 |
| Protein residues | 7378 | 3682 | 3689 |
| Ligands | 42 | 14 | 21 |
| *B* factors (Å$^2$) | | | |
| Protein | 101.83 | 9.31 | 72.59 |
| Ligand | 23.53 | 2.57 | 11.03 |
| R.m.s. deviations | | | |
| Bond lengths (Å) | 0.003 | 0.004 | 0.004 |
| Bond angles (°) | 0.597 | 0.668 | 0.659 |
| Validation | | | |
| MolProbity score | 1.25 | 1.64 | 1.38 |
| Clashscore | 4.76 | 10.15 | 5.69 |
| Poor rotamers (%) | 0.00 | 0.00 | 0.00 |
| Ramachandran plot | | | |
| Favored (%) | 98.19 | 97.44 | 97.66 |
| Allowed (%) | 1.81 | 2.56 | 2.34 |
| Disallowed (%) | 0.00 | 0.00 | 0.00 |
